# Mechanoresponsive modulation of nuclear pore complex structure and function by *O*-GlcNAc

**DOI:** 10.64898/2026.07.07.737034

**Authors:** Sunandini Chandra, Kimberly J. Morgan, William L. Chadwick, Megan C. King, C. Patrick Lusk

## Abstract

Nuclear pore complexes (NPCs) control molecular exchange across the nuclear envelope, but how they tailor their selective permeability to meet the needs of specific cell types and/or environments remains poorly understood. We demonstrate that the strength of the NPC diffusion barrier differs across cell types, is particularly stringent in cultured neurons, and correlates with the *O*-linked N-acetylglucosamine (GlcNAc) modification of nucleoporins. Using conditional tools that specifically control nucleoporin GlcNAcylation, we show that GlcNAc modulates NPC permeability. Interestingly, nucleoporin GlcNAcylation is mechanosensitive, increasing in cells plated on stiff substrates, a condition where nuclear pores dilate. Indeed, we demonstrate that increasing or decreasing GlcNAcylation dilates and constricts NPCs, respectively. Further, *O*-linked N-acetylglucosamine transferase is recruited to modify NPCs upon their acute constriction during osmotic shock. Thus, cells employ GlcNAc to modulate steady-state NPC permeability in response to mechanical inputs and to counteract critical changes to their osmotic environment.

## Introduction

Nuclear pore complexes (NPCs) are protein conduits embedded in the nuclear envelope that control molecular traffic between the nucleus and cytoplasm in all eukaryotes. Curiously, despite their essential nature, genetic variants that encode their component parts, the nucleoporins (nups), can cause several tissue-specific diseases, suggesting cell type-specific effects (Braun et al., 2026). Following on this theme, there are examples where some cell types are more sensitive than others to genetic perturbation of specific nup genes (Borlido et al., 2018; Guglielmi et al., 2024; Lupu et al., 2008; Sakuma et al., 2022; Taniguchi et al., 2025); moreover, specific nups are mislocalized or lost from NPCs in the context of neurodegenerative diseases like amyotrophic lateral sclerosis (ALS) (Chou et al., 2018; Zhang et al., 2015), observations recapitulated in cultured neuron models (Coyne et al., 2021; Coyne et al., 2020). Together, these studies support an emerging concept that there may be plasticity in the molecular composition and function of NPCs allowing them to tailor their selective permeability to the unique demands of specific cell types found within distinct tissues and environments (Cho and Hetzer, 2020; Dultz and Doye, 2025; Guglielmi et al., 2020; Lusk et al., 2025). The underlying nature of such plasticity remains ill-defined.

There are thousands of NPCs across the surface of a typical mammalian cell nucleus. These 100 MDa complexes are composed of ∼30 nups that assemble in multiples of 8, reflecting the structure’s octagonal radial symmetry (Petrovic et al., 2026). The core scaffold of the NPC structure is formed by three concentric ring assemblies: the cytoplasmic, inner and nuclear rings (Akey et al., 2022; Bui et al., 2013; Kim et al., 2018; Mosalaganti et al., 2022; Schuller et al., 2021). This scaffold directly embeds the NPC into the nuclear pore membrane through transmembrane and amphipathic helix-containing nups while also providing anchor sites for peripheral elements like the nuclear basket (Obarska-Kosinska et al., 2026; Singh et al., 2024) and cytoplasmic filaments (Bley et al., 2022; Fernandez-Martinez et al., 2016). The scaffold also delimits a central transport channel that is filled with intrinsically disordered proteins rich in phenylalanine-glycine (FG) repeats (Petrovic *et al*., 2026). This FG-rich network establishes the selective permeability of NPCs by forming a diffusion barrier that impedes passage of macromolecules greater than ∼2.6 nm and ∼40 kDa (Keminer and Peters, 1999; Mohr et al., 2009; Paine et al., 1975; Popken et al., 2015; Timney et al., 2016). Simultaneously, macromolecules with a remarkable range of molecular weights (from kDa to MDa), dimensions (up to ∼40 nm (Paci et al., 2020; Panté and Kann, 2002)) and physiochemical properties can cross this diffusion barrier when in complex with nuclear transport receptors (NTRs, karyopherins/kaps, importins, exportins, biportins (Wing and Chook, 2026)) that directly engage with the FG-repeats themselves (Bayliss et al., 2002; Bayliss et al., 2000; Hough et al., 2015; Kapinos et al., 2018; Milles et al., 2015; Port et al., 2015).

Although there is a consensus that the FG-network defines the selective permeability of NPCs, the underlying molecular mechanisms are debated. The most broadly considered models describe the FG-network in physical terms as either a biomolecular condensate with liquid (Celetti et al., 2020) or hydrogel-like properties (Frey and Görlich, 2007; Frey et al., 2006; Labokha et al., 2013; Schmidt and Görlich, 2015), or an entropic brush (Lim et al., 2006; Rout and Aitchison, 2001; Rout et al., 2000). To oversimplify, the models diverge at the ascribed importance of cohesive intra-FG interactions that are thought to be central to forming a condensate, observed *in vitro* (Frey *et al*., 2006) and *in silico* (Dekker et al., 2023). However, whether this physiochemical ability manifests *in vivo* in the presence of other abundant macromolecules in the cellular milieu is still being probed (Tetenbaum-Novatt et al., 2012; Yu et al., 2025). Moreover, how the binding of NTRs to FG-nups impacts the physical and functional state of the permeability barrier is an area of active investigation (Barbato et al., 2020; Dekker et al., 2025; Kalita et al., 2022; Kapinos *et al*., 2018; Kozai et al., 2025; Lowe et al., 2015). It is worth noting, however, that direct investigation of the FG-network in living cells remains a major technical hurdle (Yu et al., 2023; Yu *et al*., 2025). In contrast, indirect measurements of the passage of model reporter proteins through NPCs have proven valuable as a proxy to read out the central channel properties (Mohr *et al*., 2009; Popken *et al*., 2015; Timney *et al*., 2016). For example, an entropic barrier model was supported by probing the NPC diffusion barrier in yeast where even large macromolecules could cross the nuclear envelope, albeit inefficiently, suggesting a “soft” barrier (Timney *et al*., 2016). There is considerable evidence, however, for the existence and functional relevance of cohesive intra-FG interactions both *in vivo* (Hülsmann et al., 2012; Patel et al., 2007) and *in vitro* (Frey and Görlich, 2007; Frey *et al*., 2006; Schmidt and Görlich, 2015). Thus, one way to reconcile these data is to consider both models as manifesting on either end of a continuum of regimes and/or to invoke mechanisms that may toggle the behavior of the barrier between these two regimes.

There are several plausible mechanisms for how cells may tune the selective permeability of NPCs. In one example, there could be changes to the stoichiometry of nups within individual NPCs. Consistent with this idea, there are NPCs with a unique nup composition among different cell types (Cho et al., 2009; D’Angelo et al., 2012; Olsson et al., 2004; Ori et al., 2013) and even some emerging examples of individual cells with structurally different NPCs (Akey *et al*., 2022; Singh *et al*., 2024). It is worth noting, however, that these molecular changes appear primarily in the peripheral elements of the NPC including the pore membrane protein, Nup210/gp210 (D’Angelo *et al*., 2012; Olsson *et al*., 2004), or nuclear basket components (Ori *et al*., 2013; Sabinina et al., 2021; Singh *et al*., 2024). Thus, it is unclear how these molecular changes could directly impact NPC permeability. In another example, there is evidence that the NPC scaffold (and by extension, the central transport channel) may dilate and constrict (Zimmerli et al., 2021) in response to changes in nuclear envelope tension (Akey *et al*., 2022; Hoffmann et al., 2025; Mosalaganti *et al*., 2022; Schuller *et al*., 2021; Taniguchi *et al*., 2025; Zimmerli *et al*., 2021), which is itself toggled by mechanical inputs from intra and extracellular forces or from osmotic pressure (Biswas et al., 2025; Carley et al., 2021; Enyedi et al., 2016; Miroshnikova and Wickström, 2022; Morgan et al., 2025). Interestingly, there may be a spectrum of NPC dilatory states on a given nucleus, suggested to reflect local regions of nuclear envelope tension (Morgan *et al*., 2025; Mosalaganti *et al*., 2022; Schuller *et al*., 2021), at least some of which are impacted by the transmission of mechanical force through Linker of Nucleoskeleton and Cytoskeleton (LINC) complexes (Elosegui-Artola et al., 2017; Morgan *et al*., 2025). Whether NPCs of different dilatory states have unique selective permeability remains to be firmly established experimentally, but there is there is evidence that the conditions in which NPCs are the most dilated coincide with more permissive passive diffusion (Andreu et al., 2022; Hoffmann *et al*., 2025; Zimmerli *et al*., 2021), possibly facilitating import of mechanosensitive transcription factors (Elosegui-Artola *et al*., 2017). Consistent with this, the application of mechanical force on the nuclear envelope leads to an enhancement of both protein diffusion and active NTR-driven nuclear import (Andreu *et al*., 2022) with the important caveat that it is not clear whether such changes in transport kinetics are due to NPC diameter changes per se. Thus, there is a need to consider other mechanisms that may be at play.

Lastly, a particularly attractive candidate mechanism by which cells could dynamically control NPC selective permeability is through post-translational modifications like *O*-linked N-acetylglucosamine (GlcNAc)(Yang and Qian, 2017). GlcNAc is primarily placed on serines and threonines, modifying thousands of proteins including FG-nups (Hanover et al., 1987; Junod et al., 2025; Labokha *et al*., 2013; Snow et al., 1987; Zhu et al., 2016). The steady-state GlcNAcylated state of all proteins is established by an interplay between two enzymes: *O*-GlcNAc transferase (OGT) and *O*-GlcNAcase (OGA) that add and remove GlcNAc, respectively (Yang and Qian, 2017). How (or if) these enzymes are regulated to specifically modulate GlcNAcylation of nups is unknown, but there is evidence that globally reducing GlcNAcylation decreases the stringency of the NPC diffusion barrier while increasing NTR-driven transport kinetics (Yoo and Mitchison, 2021). These changes to the NPC may also explain why mRNA export frequency is higher when nups are GlcNAcylated (Junod *et al*., 2025). One caveat to these studies is that nup GlcNAcylation was modulated using drugs that globally alter the GlcNAcylation of all proteins, leaving open the possibility that the observed impact on NPCs is indirect. However, *in vitro* evidence supports that the introduction of the hydrophilic GlcNAc modification into the hydrophobic FG-network is sufficient to dampen cohesive FG-FG interactions required to establish a robust diffusion barrier (Labokha *et al*., 2013). The coherence between the *in vitro* and *in vivo* data strongly motivates the development of approaches to examine the consequences of GlcNAcylation on FG-nup behavior (Yu *et al*., 2025) and more selective tools to directly inhibit or enhance nup GlcNAcylation in living cells.

Here, by leveraging a series of fluorescent reporters with a spectrum of molecular weights (Timney *et al*., 2016), we uncover that diffusion across NPCs varies considerably across cell types. We correlate these diffusion barrier differences to nup GlcNAcylation levels, which are particularly low in neurons where the diffusion barrier is most stringent. Interestingly, using genetically-encoded tools we demonstrate that increasing or decreasing nup GlcNAcylation drives dilation or constriction of NPCs, respectively, supporting that GlcNAcylation of nups can modulate NPC diameter. Indeed, GlcNAcylation of nups is responsive to mechanical inputs and is most elevated when cells are cultured in stiff mechanical environments coincident with NPC dilation. Further, OGT is rapidly deployed to the NPC in response to acute NPC constriction driven by osmotic stress, supporting that nup GlcNAc dynamically modulates the selective permeability of NPCs in response to both acute and chronic mechanical inputs.

## Results

### Model neuronal cell lines have a stringent NPC diffusion barrier

To test whether unique cell types have distinct NPC diffusion barrier properties we first compared HeLa cells to differentiated, “neuron-like” SH-SY5Y cells (Biedler et al., 1978). Note that although differentiated (d) SH-SY5Y cells have fewer NPCs than HeLa cells, they are smaller such that the density of NPCs between these cell types and their karyoplasmic ratios are comparable (Morgan *et al*., 2025). We took advantage of previously described EGFP-based reporters that contain from 1 to 6 Protein A (PrA) moieties and range from 27-67 kDa (Fig. 1A, S1A (Andreu *et al*., 2022; Timney *et al*., 2016)); importantly, the shape of the reporters was found to have little effect on their diffusivity (Timney *et al*., 2016). To measure and quantify passive diffusion across the nuclear envelope, we photobleached the fluorescence in the cytoplasm and measured the loss of nuclear fluorescence over time (Fig. 1B). Quantitative differences in the rate of nuclear fluorescence loss of the reporters between HeLa and dSH-SY5Y cells was apparent in single cell traces, which became slower as reporter size increased as expected (Fig. 1C and D). Plotting the rate of nuclear fluorescence loss from several cells from multiple experimental replicates further reinforced this assessment (Fig. 1E). Interestingly, although the larger 47-67 kDa reporters effluxed from the nucleus with similar kinetics between the two cell types, there were obvious differences between the behavior of the smaller constructs. For example, the 27 kDa reporter effluxed at about half the rate in the neuronal cell type compared to HeLa cells (Fig. 1E). Together, these data support that although NPCs in both cell types impose a size-dependent diffusion barrier, there is an overall more stringent barrier in the neuron model that is most apparent at relatively lower molecular weights.

**Figure 1.**
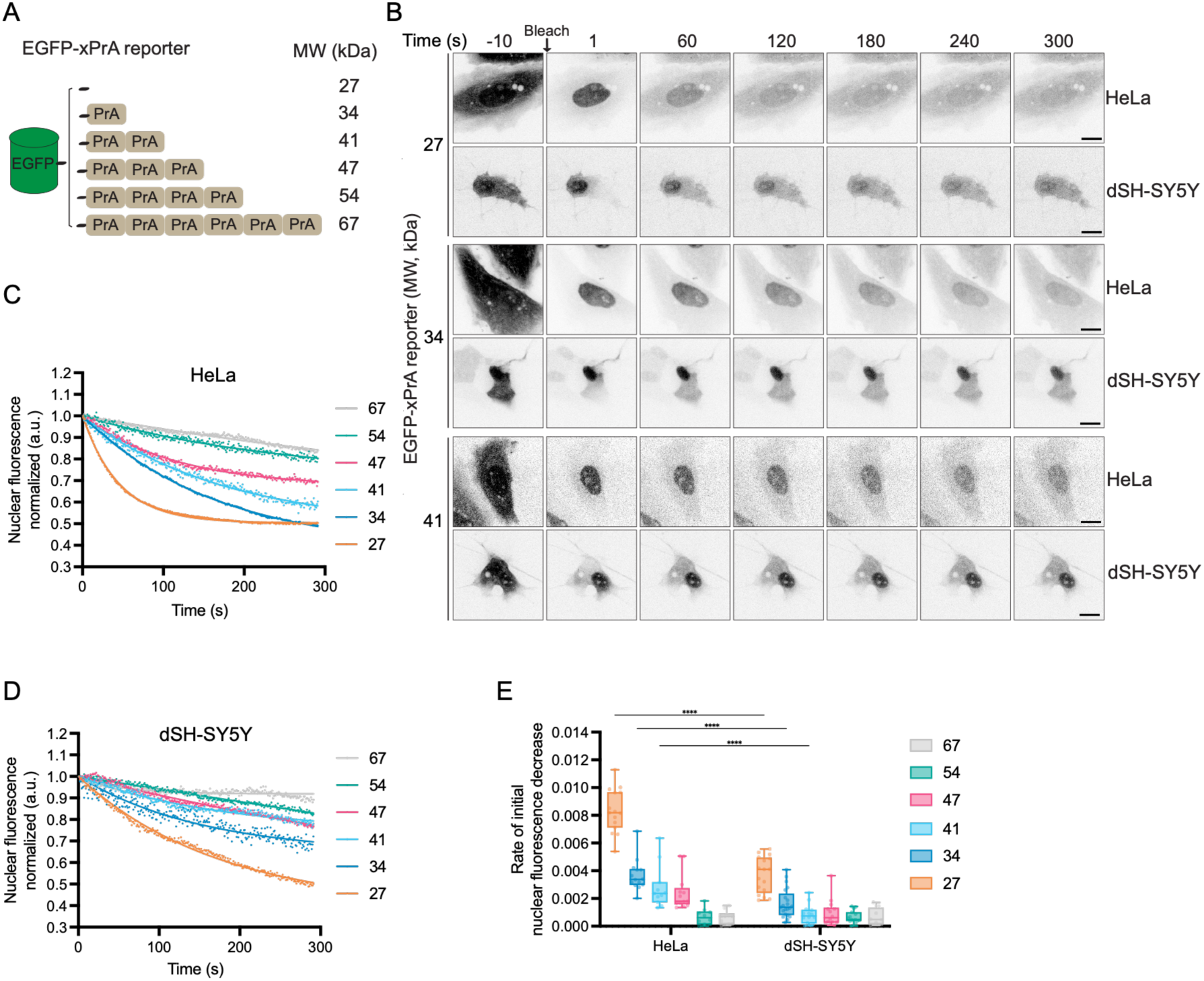
The passive diffusion barrier is more stringent in differentiated neuron-like dSH-SY5Y cells than HeLa cells. (A) Schematic of EGFP-xPrA reporters used in this study. (B) Confocal fluorescence micrographs of HeLa or dSH-SY5Y cells transiently expressing 27, 34, or 41 kDa EGFP-xPrA reporters. Cytoplasmic reporter fluorescence was bleached, and nuclear fluorescence was monitored at the indicated time points (seconds). Scale bar, 20 μm. (C, D) Plots of the nuclear fluorescence of the indicated EGFP-xPrA reporter from single HeLa (C) or dSH-SY5Y (D) cells after cytoplasmic fluorescence was bleached (Time=0 is 20 s after bleaching). Data were fitted using a one-phase exponential decay model by nonlinear least-squares regression. (E) Rate of decrease of nuclear fluorescence from all replicates of each reporter and cell line derived from the slope of a straight line plotted from initial 40 s of the 300 s time course. Data represented as box and whiskers plot, box boundaries represent interquartile range, whiskers extend from minimum to maximum data point. Each data point represents a cell. For each sample n=10-21 cells from 3 biological replicates and the statistical test is two-way ANOVA with Tukey’s method for multiple comparisons. ****, p<0.0001.

To evaluate whether a stringent NPC diffusion barrier may be a property of neurons more generally, we examined the diffusion rates of the 27, 34 and 41 kDa GFP constructs in human induced pluripotent stem cell (iPSC)-derived integrated, inducible, and isogenic lower motor neurons (i^3^LMNs (Fernandopulle et al., 2018); Fig. 2A). Comparing again the single cell traces of the 27, 34 and 41 kDa GFP reporters, there was a clear trend in which the nuclear fluorescence loss was slower in the i^3^LMNs compared to their undifferentiated iPSC counterparts (Fig. 2B, C) with mean rates of nuclear fluorescence loss significantly slower in the i^3^LMNs (Fig. 2D). Indeed, the 41 kDa reporter did not appreciably leave the nucleus of this example i^3^LMN over the 300 s time course. Thus, like the dSH-SY5Y cells, i^3^LMNs impose a stringent barrier to macromolecular diffusion across their NPCs. Indeed, i^3^LMNs showed the slowest diffusion rate of the 34 kDa reporter across all cell lines that we tested, including A549, HCT116 and U2OS (Fig. 2E).

**Figure 2.**
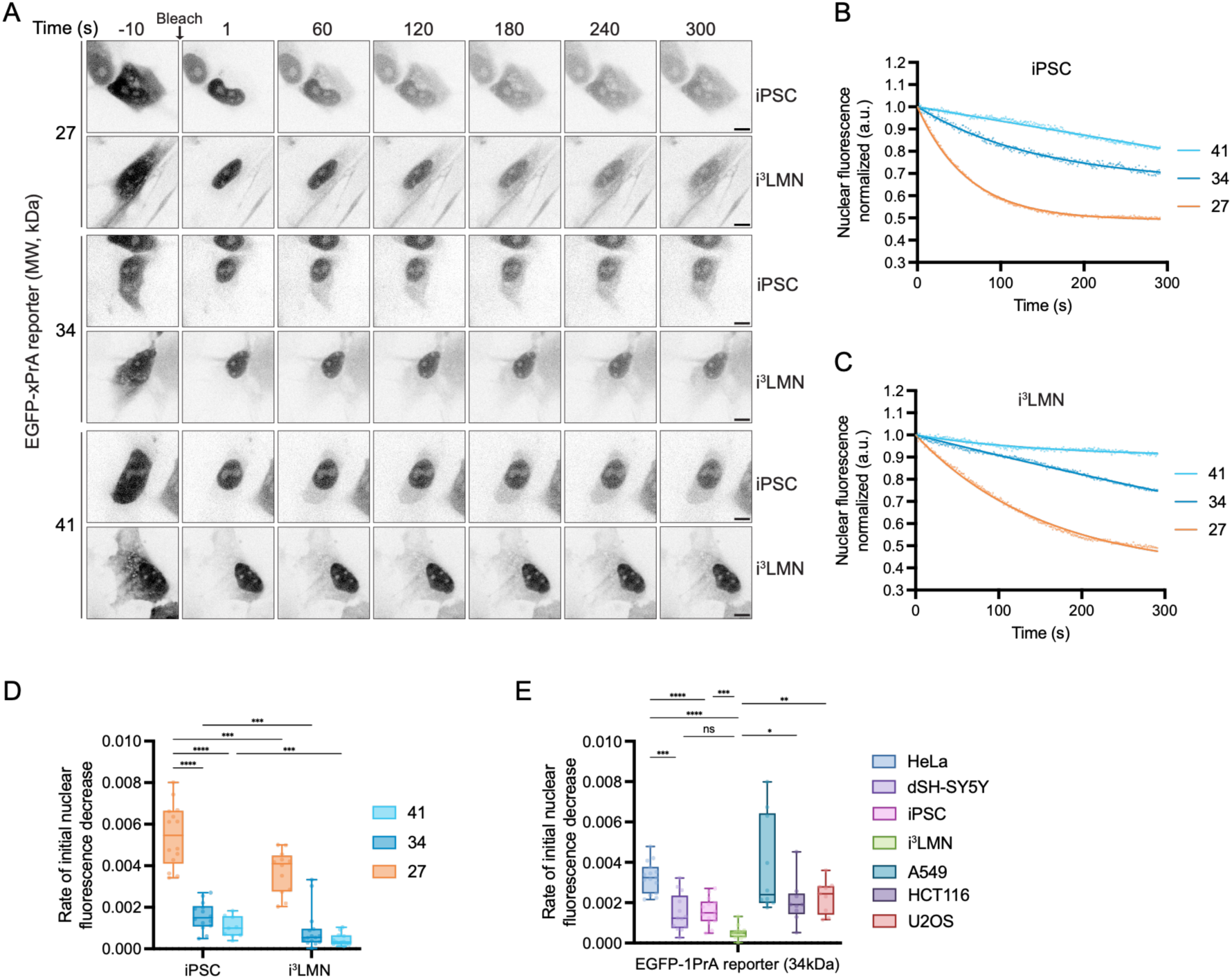
The passive diffusion barrier is more stringent in i^3^LMNs than in the parental iPSCs. (A) Confocal fluorescence micrographs of iPSCs or i^3^LMNs transiently expressing 27, 34, or 41 kDa EGFP-xPrA reporters. (B, C) Plots of the nuclear fluorescence of the indicated EGFP-xPrA reporter from single iPSC (B) or i^3^LMN (C) cells after cytoplasmic fluorescence was bleached (Time=0 is 20 s after bleaching). Data were fitted using a one-phase exponential decay model by nonlinear least-squares regression. (D) Rate of decrease of nuclear fluorescence from all replicates of each reporter and cell line derived from the slope of a straight line plotted from the initial 40 s of the 300 s time course. Data represented as box and whiskers plot, box boundaries represent interquartile range, whiskers extend from minimum to maximum data point. Each data point represents a cell. For each sample n=12-15 cells from 2 biological replicates and the statistical test is two-way ANOVA with Tukey’s method for multiple comparisons. ***, p=0.0003; ****, p<0.0001. (E) Rate of decrease of nuclear fluorescence as in D, shown for the indicated cell lines expressing 34 kDa EGFP-1PrA reporter. Statistical test is Brown-Forsythe and Welch’s ANOVA with Dunnett’s T3 method for multiple comparisons. ns, p=0.0513; *, p=0.0439, **, p=0.0019; ***, p=0.003 for HeLa vs dSH-SY5Y and p=0.009 for iPSC vs i^3^LMN; ****, p<0.0001

### GlcNAcylation of nups modulates the NPC diffusion barrier

Recent work has implicated nup GlcNAcylation as a mechanism for modulating the relative stringency of the NPC diffusion barrier (Yoo and Mitchison, 2021). We therefore tested whether differences in nup GlcNAcylation could explain the diffusion barrier properties of the cultured neurons. We first examined nup GlcNAcylation with a commonly employed monoclonal antibody (RL2) that recognizes GlcNAc on proteins with a preference for GlcNAcylated nups (Snow *et al*., 1987). To confirm that this antibody recognized the most heavily GlcNAcylated nups (Ma et al., 2021), we depleted the transcripts for *NUP214*, *NUP153*, *NUP62* and *NUP98* by siRNA and observed the specific loss of the appropriate RL2-reactive bands on immunoblots of total protein extracts (Fig. S1B). Armed with this knowledge, we directly compared RL2 immunoblots of HeLa and dSH-SY5Y protein preparations. Strikingly, we observed less RL2 signal in the dSH-SY5Y samples (Fig. 3A). By performing densitometry across the entire gel lanes, we measured 60% less signal in the nup-specific bands of the dSH-SY5Y samples (Fig. 3A). The lower nup GlcNAc levels in the dSH-SY5Y samples were not the result of an alteration in relative abundance of NPCs, as assessed by Nup62 levels, nor changes to the OGA or OGT enzymes themselves (Fig. 3A). We further confirmed a dearth of nup GlcNAcylation in dSH-SY5Y cells *in vivo* using a metabolic labeling approach where GlcNAc is replaced with a GlcNAz moiety that can be dye-labeled using click chemistry (Yoo and Mitchison, 2021). As shown in Fig. 3B, the fluorophore-GlcNAz signal at the nuclear rim visible in HeLa cells is notably absent in dSH-SY5Y cells, which is also obvious when quantified by relating the intensity of the nuclear rim GlcNAz signal to mAb414-labeled NPCs (mAb414 recognizes FxFG-nups (Davis and Blobel, 1986); Fig. 3C). We further noted a ∼30% reduction in RL2 signal in i^3^LMNs when compared to their iPSC parents after normalizing for a modest reduction in Nup93 protein levels (Fig. 3D). Thus, the low nup GlcNAcylation in these cultured neuron models provides a plausible explanation for their more stringent NPC diffusion barriers.

**Figure 3.**
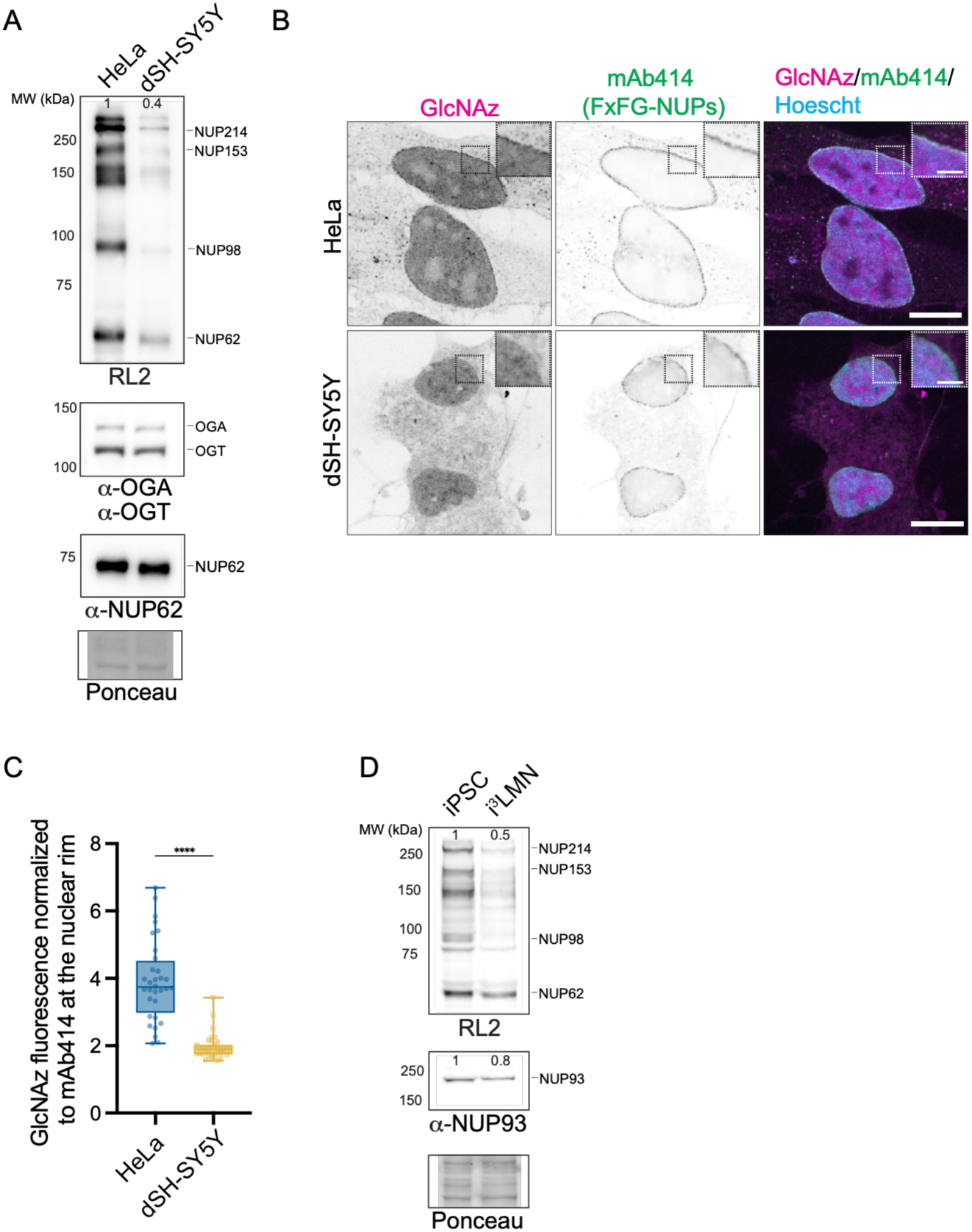
Nup GlcNAcylation is low in cultured neurons. (A) Immunoblots (one of three independent biological replicates shown) of equivalent protein loads from whole cell extracts of HeLa and dSH-SY5Y cells. GlcNAc conjugates detected by the RL2 antibody with specific nup positions at right, and the indicated proteins with specific antibodies. Numbers on the left indicate positions of molecular weight (MW) standards in kDa. Numbers on top of each lane indicate relative amount of RL2 band intensity in dSH-SY5Y compared to HeLa cells. To confirm equal protein loading, a portion of the blot stained with Ponceau S is shown. (B) Confocal fluorescence micrographs of HeLa and dSH-SY5Y cells metabolically labeled with GlcNAz, detected by CuAAC labeling with CF640R. FxFG-Nups detected with the mAb414 antibody and Alexa Fluor 488-conjugated secondary antibody. DNA stained with Hoechst. Scale bar 10 μm. Inset scale bar 5 μm. (C) The fluorescence of CF640R-labeled GlcNAz was normalized to the mAb414 signal along an ROI encompassing the nuclear rim of n=32 cells from 2 independent experiments. Data represented as box and whiskers plot, box boundaries represent interquartile range, whiskers extend from minimum to maximum data point, statistical test is an unpaired t-test with Welch’s correction. ****, p<0.0001. (D) Immunoblots as in A (one of two representative independent biological replicates shown) of proteins derived from whole cell extracts of iPSCs and i^3^LMNs. Numbers on top of each lane indicate relative amount of RL2 or NUP93 band intensity in i^3^LMNs compared to iPSCs.

### Engineered modification of nups alters NPC diffusion barrier properties

To provide evidence that the relatively high and low levels of nup GlcNAc in HeLa and dSH-SY5Y cells explained the diffusion barrier properties of these cell types, we employed OSMI-4 and Thiamet G (TMG) that inhibit the OGT and OGA enzymes, respectively. Treatment of HeLa cells with OSMI-4 markedly reduced nup GlcNAcylation as expected (Fig. S2A). Further, we observed a decrease in the relative rate of diffusion of a EGFP reporter across the nuclear envelope (Fig. S2B). Conversely, treatment of dSH-SY5Y cells with TMG increased nup GlcNAc levels (Fig. S2A) and consequently elevated the passive diffusion rate of the reporter (Fig. S2B).

As thousands of proteins are modified by GlcNAc (Ma *et al*., 2021), the impacts of OSMI-4 and TMG treatment on nucleocytoplasmic transport may be indirect. Therefore, to explore the sufficiency of nup GlcNAc modification to modulate the diffusion barrier of NPCs in different cell types, we developed tools to conditionally alter nup GlcNAcylation without modifying whole proteome GlcNAcylation patterns. We generated constructs where OGT or OGA were fused to a HaloTag and GFP binding protein (GBP), which were then transfected into U2OS cells CRISPR-edited to express a NUP96-mEGFP fusion from its endogenous locus. The expression of Halo-GBP-OGT was induced with doxycycline and, as designed, Halo-GBP-OGT was efficiently recruited to the nuclear periphery (Fig. 4A). As a comparison, the endogenous enzyme is distributed throughout the cell, with a bias for the nucleus (Fig. S2C). Nup GlcNAcylation was assessed by RL2 staining, which was clearly elevated along the nuclear rim in Halo-GBP-OGT expressing cells (Fig. 4A). Further, as RL2 also recognizes other GlcNAcylated species, we performed line profiles that captured segments of the cytoplasm and nucleoplasm in the presence or absence of Halo-GBP-OGT: the RL2 signal specific to expression of Halo-GBP-OGT was at the nuclear envelope (Fig. 4B, C). Importantly also, the Halo-GBP-OGA or the catalytically dead Halo-GBP-OGA(D174N)(Cetinbaş et al., 2006) were both recruited efficiently to NUP96-GFP (the endogenous enzyme is distributed throughout the cell, Fig. S2C), however, only Halo-GBP-OGA reduced fluorescence of RL2 along the nuclear rim (Fig. 4D-F).

**Figure 4.**
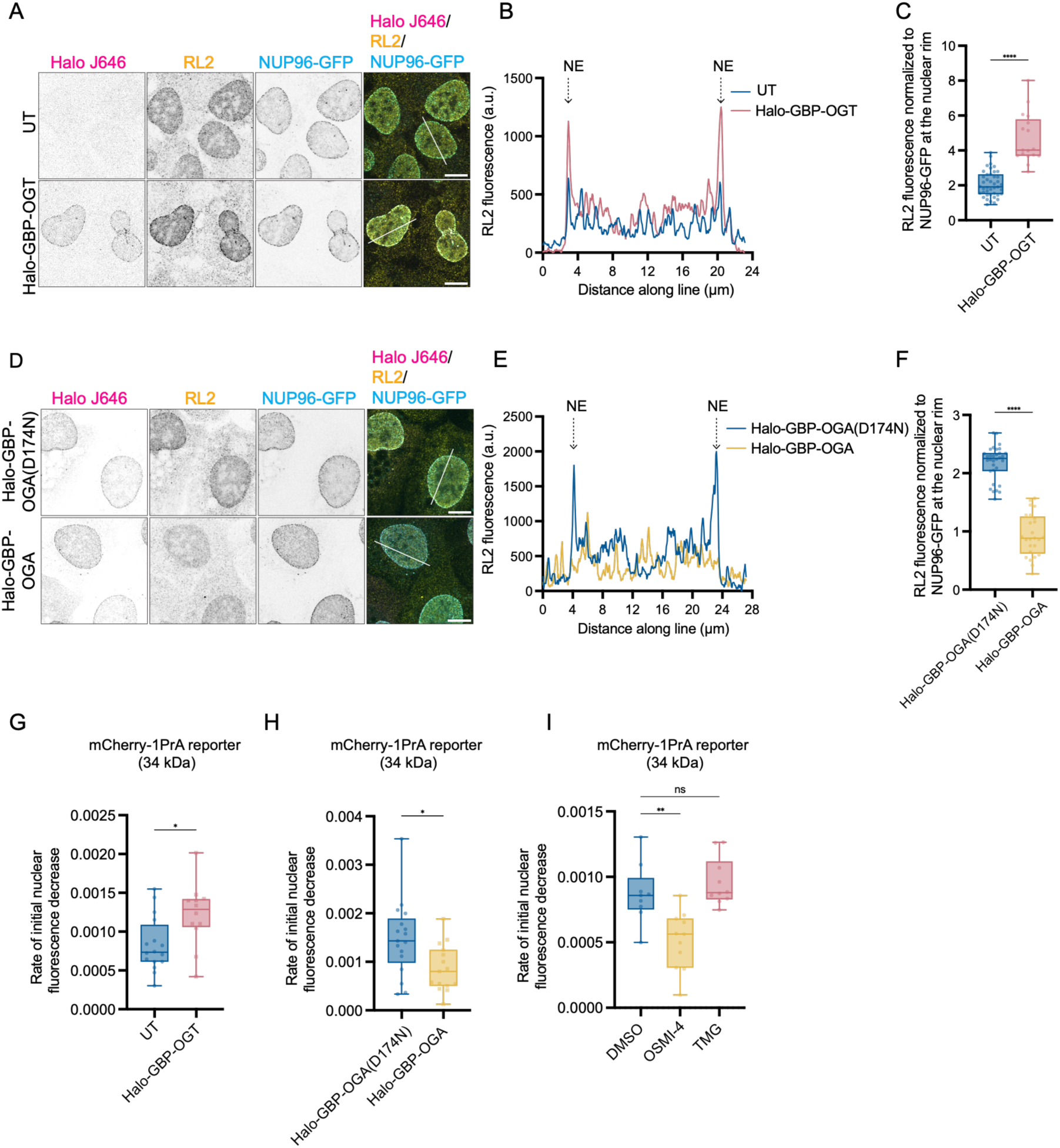
Direct modulation of nup GlcNAcylation impacts NPC passive diffusion rates. (A) Confocal immunofluorescence images of U2OS-CRISPR-NUP96-mEGFP cells with or without (untransfected/UT) the expression of Halo-GBP-OGT. Cells stained with the J646 Halo ligand and the RL2 antibody followed by Alexa Fluor 594-conjugated secondary antibody, scale bar 10 μm. (B) Plot of the fluorescence of the RL2 signal along indicated lines in the right most panels of (A). Blue lines, UT; pink line, expressing Halo-GBP-OGT. Position along the nuclear envelope (NE) assessed by NUP96-GFP position (not shown). (C) Plot of the RL2 signal normalized to NUP96-GFP fluorescence in UT cells or those with Halo-GBP-OGT. Data represented as box and whiskers plot, box boundaries represent interquartile range, whiskers extend from minimum to maximum data point. n>17 cells from 2 independent biological replicates, statistical test is two-tailed Mann Whitney. ****, p<0.0001. (D-F) As in (A-C) but comparing cells expressing catalytically-active Halo-GBP-OGA to those expressing catalytically-inactive Halo-GBP-OGA(D174N). (G-I) Rate of initial nuclear fluorescence loss of 34 kDa mCherry-1PrA reporter plotted as in Fig. 1E. (G) U2OS-CRISPR-NUP96-mEGFP cells UT or expressing Halo-GBP-OGT. n>14 cells from n=2 independent experiments. Statistical test is two-tailed Mann Whitney. *, p=0.0282. (H) U2OS-CRISPR-NUP96-mEGFP cells expressing catalytically-active Halo-GBP-OGA or catalytically-inactive Halo-GBP-OGA(D174N). n>16 cells from 3 biological replicates, statistical test is two-tailed Mann Whitney. *, p=0.0158. (I) U2OS cells after 24 h treatment with DMSO, OSMI-4 or TMG n=10 cells from two independent biological replicates. Statistical test is Brown-Forsythe and Welch’s ANOVA with Dunnett’s T3 method for multiple comparisons. ns, p=0.5990; **, p=0.0048.

With these tools in hand, we tested how specific reduction or elevation of nup GlcNAcylation impacted the diffusion rates of a mCherry-1PrA reporter across the nuclear envelope. Consistent with the interpretation that the GlcNAcylation of nups controls the diffusion barrier properties of NPCs, we observed a faster diffusion rate of the mCherry-1PrA reporter in Halo-GBP-OGT-expressing cells compared to the untransfected control (Fig. 4G). Reciprocally, the diffusion rate of mCherry-1PrA was slower in the Halo-GBP-OGA-expressing cells when compared to those expressing the catalytically dead Halo-GBP-OGA(D174N)(Fig. 4H). Last, we compared the pharmacological approach with the expression of Halo-GBP-OGT and Halo-GBP-OGA and found that they had near analogous effects on the diffusion barrier of the NUP96-GFP U2OS cells: OGT inhibition with OSMI-4 (Fig. S2D) increased the stringency of the NPC diffusion barrier, mimicking the targeting of the OGA enzyme (compare Fig. 4I and Fig. 4G). However, OGA inhibition with TMG (Fig. S2D) does not lead to a significant effect on mCherry-1PrA diffusion (Fig. 4I). This suggests that driving OGT to the NPC to specifically enhance nup GlcNAcylation may be more effective at decreasing the stringency of the NPC diffusion barrier than disrupting OGA, which is required for GlcNAc removal. Taken together, our data demonstrate that local modulation of GlcNAcylation at the NPC can modulate the NPC barrier while also providing additional confidence that pharmacological modulation of OGT/OGA can also alter the NPC diffusion barrier properties even in the context of global changes to GlcNAc.

### GlcNAcylation contributes to NPC dilation state

To investigate the mechanism(s) by which GlcNAc contributes to defining the stringency of the NPC diffusion barrier, we performed pan-Expansion Microscopy (pan-ExM; (M’Saad and Bewersdorf, 2020)) on iPSCs treated with OSMI-4 or TMG. Samples were expanded isotropically ∼16 fold, proteins were stained with a “pan-stain” (a fluorophore conjugated N-hydroxysuccinimide (NHS) ester), SYTOX (to visualize DNA), and RL2 (Fig. S3A, B). As we previously reported (Morgan *et al*., 2025), the pan-stain allowed comprehensive visualization of the thousands of NPCs across the entire nucleus. Further, the cytoplasmic ring (CR), inner ring (IR, with FG-nups) and nuclear ring (NR) of the NPCs can be visualized in side views (Fig. 5A). Consistent with other super-resolution microscopy studies, the RL2 signal is concentrated at the IR (Junod *et al*., 2025; Yoo and Mitchison, 2021), indicating that the majority of nup GlcNAcylation is found within the FG-network. This signal was reduced by ∼50% after treatment of cells with OSMI-4, while we observed a doubling in fluorescence in the presence of TMG. Note that even in this maximal nup-GlcNAc scenario, the majority of the RL2 signal remained co-localized with the inner ring (Fig. 5A, B).

**Figure 5.**
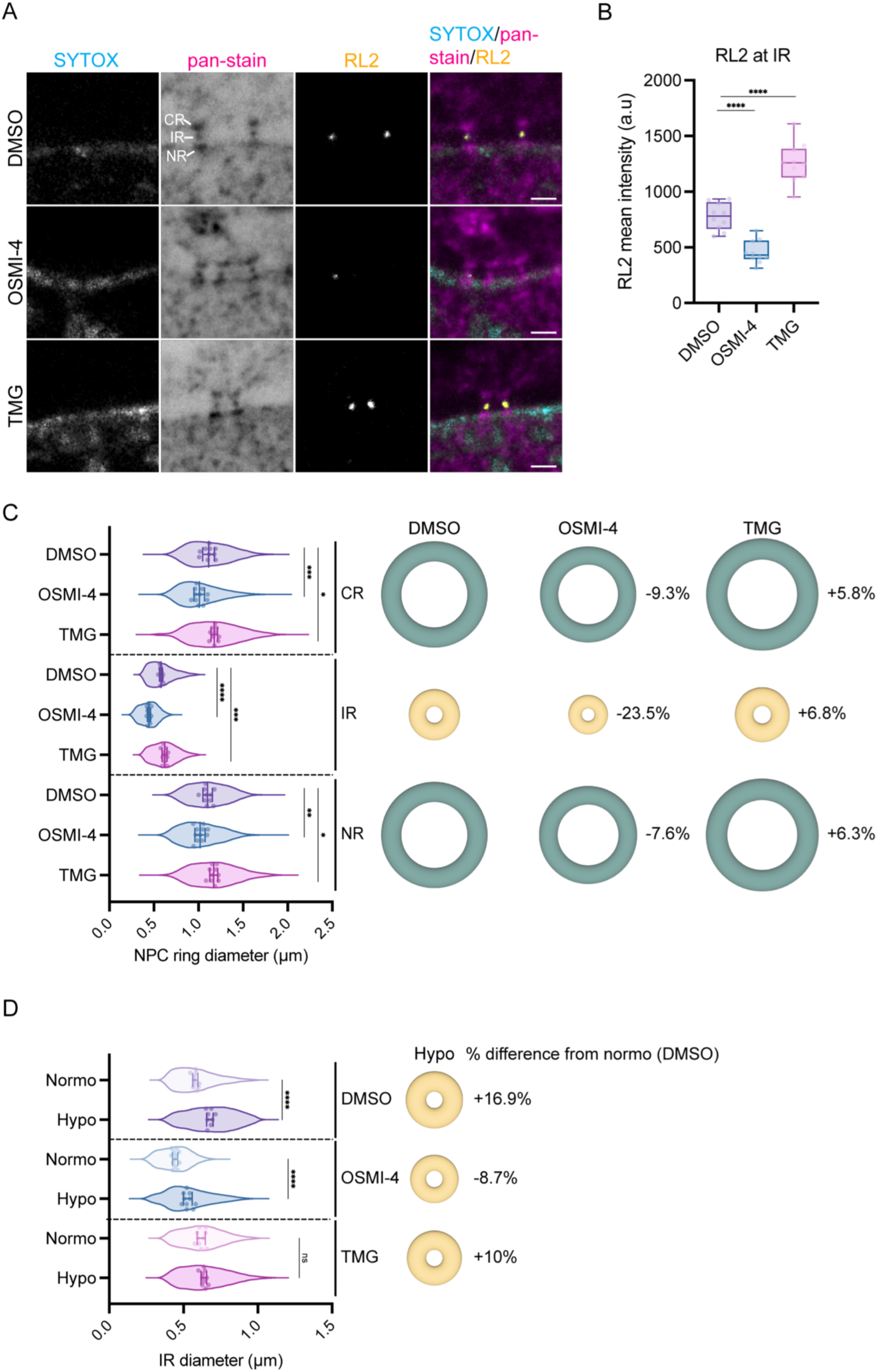
Nup GlcNAcylation contributes to NPC dilation. (A) Confocal fluorescence images of the nuclear envelope of iPSCs treated with DMSO, OSMI-4 or TMG for 24 h and subjected to pan-Expansion microscopy. SYTOX (DNA), pan-stain (NHS-Ester-CF568), and RL2 (ATTO647N) channels with merge shown. The NPC architecture is visualized in the pan-stain. CR=cytoplasmic ring, IR=inner ring, NR=nuclear ring. Scale bar of the expanded sample, 2 µm. (B) Plot of the mean fluorescence intensity in a.u. of the RL2 signal at the inner ring (IR) from iPSCs treated as indicated. Data represented as box and whiskers plot, box boundaries represent interquartile range, whiskers extend from minimum to maximum data point. Symbols denote individual cell means. n=10 cells. Ordinary one-way ANOVA with Dunnet’s method for multiple comparisons. ****, p<0.0001. (C) Measurement of the NPC ring diameter at the CR, IR and NR from expanded, pan-stained iPSCs treated as indicated. Symbols denote individual cell means. Line and error bars are the overall mean ± SD. n=109,172 NPC rings total in n=10-12 cells from one expansion experiment. Ordinary one-way ANOVA with Dunnet’s method for multiple comparisons. *, p=0.0283; ***, p=0.0003; ***, p=0.0005; ****, p<0.0001; *, p=0.0122; **, p=0.0017. The diameter of the indicated NPC ring and drug treatment is plotted and schematized at right. The numbers in the ring diagrams represent the percent change from the DMSO condition. (D) The diameter of the NPC IR in the indicated osmotic conditions (left) and drug treatment (right) is plotted and schematized at the right. The numbers in the ring diagrams represent the % change from the normo osmotic DMSO condition. n=73,939 IRs total in 8-12 cells from one expansion experiment. Mixed-effects model with Dunnet’s method for multiple comparisons. ****, p<0.0001; ns, p=0.0901.

Recent evidence supports that the application of mechanical force on the nucleus increases nuclear transport rates and lowers the stringency of the NPC diffusion barrier, perhaps by dilating NPCs (Andreu *et al*., 2022). We therefore used pan-ExM to examine whether altering nup GlcNAcylation was sufficient to influence NPC diameter. For this analysis, we used a previously developed machine learning-based algorithm (Morgan *et al*., 2025) to first segment and then measure the diameters of each of the three NPC rings of all NPCs in iPSCs treated with carrier alone (DMSO), OSMI-4, or TMG. Strikingly, depletion of nup GlcNAcylation led to a significant 23.5% decrease in the diameter of the IR and more modest 7.6 to 9.3% diameter reductions of the NR and CR, respectively, consistent with the bulk of nup GlcNAcylation residing in the IR (Fig. 5A, C). Elevating GlcNAc increased the diameters of all three rings by ∼6% (Fig. 5C). Note that we confirmed that neither the OSMI-4 nor TMG treatments impacted expansion of other cellular structures like mitochondria or centrosomes, suggesting a specific impact on NPCs (Fig. S4A, B, E).

One potential caveat to the conclusion that nup GlcNAcylation is sufficient to modulate NPC dilation is the possibility that the pharmacological approach influences the tension state of the nuclear envelope. We therefore took advantage of prior work in which a hypo-osmotic shock was tied to increased nuclear envelope tension (Finan et al., 2009; Hoffmann *et al*., 2025; Lemière et al., 2022; Mitchison, 2019). Indeed, after a 5 minute hypo-osmotic shock, we measured a 17% increase in the diameter of the IR (Fig. 5D, 17%, DMSO). To evaluate whether the presence of nup GlcNAc impacted this diameter change, we pre-treated cells with OSMI-4 to reduce nup GlcNAcylation prior to the hypo-osmotic shock. Strikingly, in this scenario the IR failed to fully dilate arguing that a poorly GlcNAcylated FG network antagonizes NPC dilation even in the context of elevated nuclear envelope tension (Fig. 5D). TMG treatment had little effect likely reflecting that iPSCs have robust GlcNAcylation at baseline (Fig. 3D). Taken together, these findings support that raising or lowering nup GlcNAcylation may be sufficient to increase or decrease NPC dilation, respectively, with a collaboration between nup GlcNAcylation and nuclear envelope tension being required to fully dilate the NPC.

### Nup GlcNAcylation is responsive to substrate mechanics

The observation that nup GlcNAcylation can impact NPC dilation prompted an investigation into the recently described sensitivity of NPC permeability to mechanical inputs (Andreu *et al*., 2022; Elosegui-Artola *et al*., 2017). Specifically, the diffusion barrier was shown to be more stringent to small 27-41 kDa reporters when cells were grown on soft substrates, implying that NPCs are found in a more constricted state due to the likely reduction in nuclear envelope tension under these conditions. Indeed, we recapitulated these findings by measuring the rate of NPC diffusion of a 34 kDa mCherry reporter (that diffuses comparably to the 41 kDa EGFP-2PrA construct, Fig. S2E) in U2OS and A569 cells grown on either a “soft” 3 kPa or “stiff” 1.5 MPa substrate. Consistent with published work (Andreu *et al*., 2022), the diffusion rates were relatively slower on the softer substrates in both cell lines (Fig. 6A).

**Figure 6.**
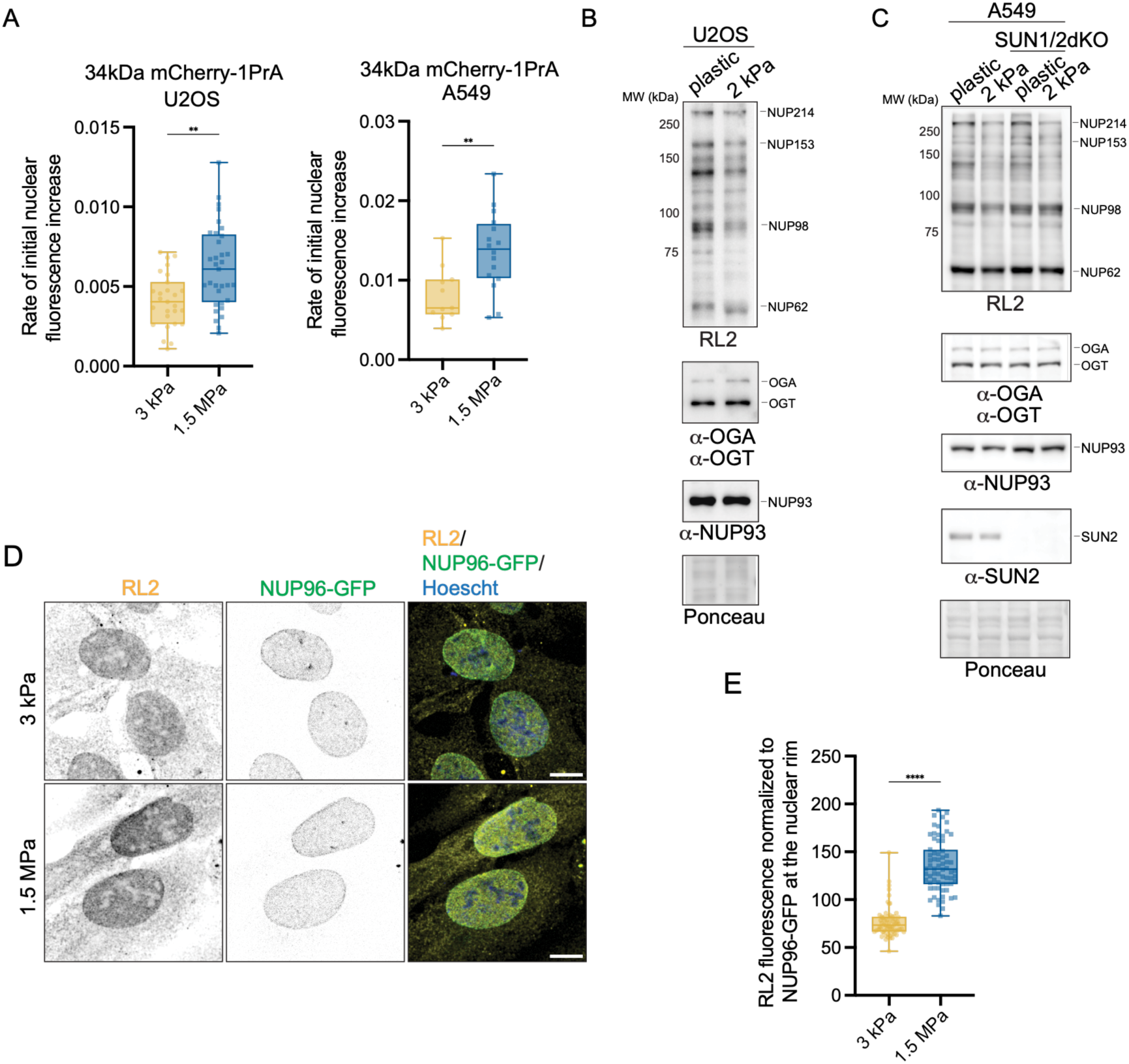
Nup GlcNAcylation is mechanosensitive. (A) Rate of initial nuclear fluorescence increase of 34 kDa mCherry-1PrA reporter expressed in U2OS and A549 cells plated on PDMS substrates prepared to be soft (3 kPa) or stiff (1.5 MPa). Plotted as in Fig. 1E. Two-tailed Mann Whitney test performed. n>27 from 2 independent experiments for each condition**, p=0.0013 for U2OS and n>11 from one experiment, p=0.0047 for A549. (B) Immunoblots from protein extracts of U2OS cells plated on soft Matrigen hydrogels (2 kPa) or plastic (GPa) substrates, probed with RL2, OGA, OGT and NUP93 antibodies. Numbers on the left indicate positions of molecular weight (MW) standards in kDa. Ponceau S stain shown to assess relative protein loading. Representative immunoblot shown from 3 biological replicates. (C) Immunoblots (representative of two biological replicates) of protein extracts as in (B) but from A549 and A549-SUN1/2dKO cells. (D) Confocal fluorescence micrographs of U2OS-CRISPR-NUP96-mEGFP cells plated on soft (3 kPa) or stiff (1.5 MPa) PDMS substrates for 48 h and stained with RL2. RL2 labeling (Alexa Fluor 647), NUP96-mEGFP and merge (with Hoechst) shown. Scale bar 10 μm. (E) Plot of the RL2 signal normalized to NUP96-GFP fluorescence in cells plated on 3 kPa or 1.5 MPa PDMS substrates. n>64 cells from 2 independent replicates for each condition. Two-tailed unpaired t-test with Welch’s correction performed. ****, p<0.0001.

To examine whether nup GlcNAcylation could explain the mechanosensitivity of the NPC’s diffusion barrier, we tested whether nup GlcNAcylation was altered in U2OS (and A549) cells grown on substrates that were relatively soft (2 kPa) or stiff (plastic/GPa). Remarkably, we observed a higher RL2 signal when probing protein extracts derived from both the U2OS and A549 cell lines plated on plastic (Fig. 6B, C). Interestingly, the higher levels of RL2 immunogenicity in the A549 cells plated on the stiff substrate was not dependent on LINC complexes as we observed an analogous trend in A549 cell lines lacking both SUN1 and SUN2 (Fig. 6C). Further, this effect was not due to any obvious changes to the levels of OGT or OGA, nor to levels of a representative nup, NUP93 (Fig. 6B, C). To further account for any potential mechanosensitive effect on NPC number, we also directly related the RL2 signal at the nuclear envelope to the fluorescence of endogenously expressed NUP96-GFP in U2OS cells grown on 3 kPa or 1.5 MPa substrates. This analysis revealed that while NUP96-GFP distribution and fluorescence intensities were comparable between the two conditions, the RL2 signal was significantly higher at the nuclear envelope on the stiffer, 1.5 MPa substrates (Fig. 6D, E). Thus, nup GlcNAcylation is responsive to mechanical inputs.

### Rapid changes to nup GlcNAcylation counteracts NPC constriction in response to hyper-osmotic shock

In considering the potential for functional crosstalk between nup GlcNAcylation, NPC dilation, and nup GlcNAc mechanosensitivity, we next examined the sufficiency of perturbing NPC dilation to alter nup GlcNAcylation. To date, only hyper- or hypo-osmotic shock has been shown to critically constrict or dilate NPCs in both yeast and mammalian cells, respectively (Hoffmann *et al*., 2025; Morgan *et al*., 2025; Zimmerli *et al*., 2021). We therefore incubated U2OS cells expressing endogenous NUP96-GFP for 30 minutes in either hyper or hypo-osmotic conditions and monitored nup GlcNAc with RL2 staining. We observed a modest decrease in the RL2 signal in the hypo-osmotic condition when normalized to the dilution of the NUP96-GFP that occurred as the nuclear surface area increased (Fig. 7A, B). However, in this case we could not be definitive as to whether this decrease was specific for the RL2 at the nuclear envelope as the RL2 at the nuclear rim was indistinguishable from that within the nucleus (insets, Fig. 7A). Far more striking was the clear enhancement of the RL2 signal along the nuclear periphery in cells subjected to the hyper-osmotic shock (Fig. 7A). Once normalized to the NUP96-GFP fluorescence that is locally elevated due to the loss of nuclear volume, we measured a robust (23.8%) increase in RL2 staining intensity, indicating an increase in nup GlcNAcylation under these conditions (Fig. 7B).

**Figure 7.**
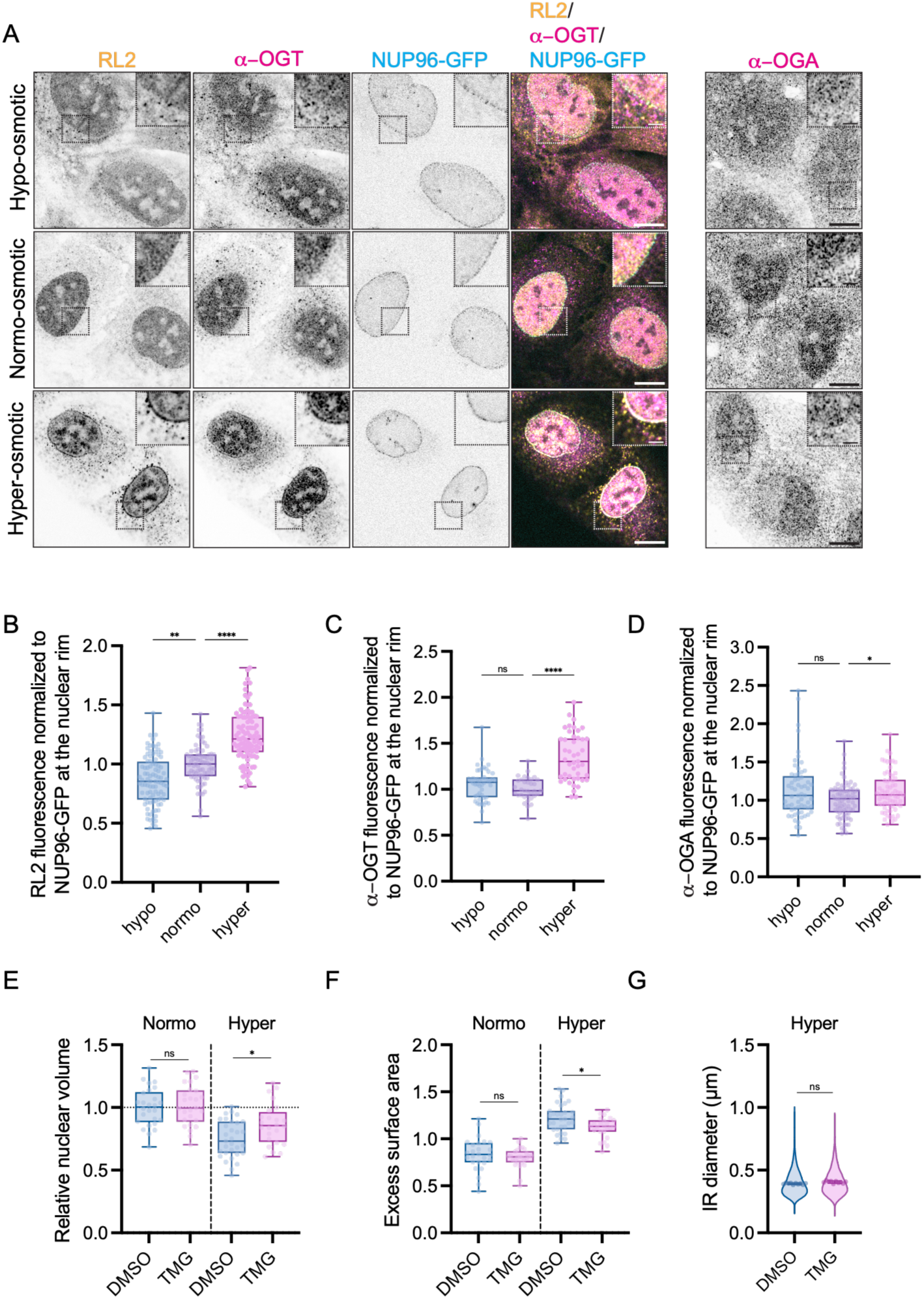
OGT drives nup GlcNAcylation in response to hyperosmotic shock. (A) U2OS-CRISPR-NUP96-mEGFP cells were exposed to normo-osmotic media, hypo-(0.5X media), or hyper- (media + 400 mM sorbitol) osmotic conditions for 30 min, fixed and prepared for immunofluorescence staining with RL2 and anti-OGT (or anti-OGA) antibodies. Confocal fluorescence micrographs of the RL2, anti-OGT, NUP96-GFP signal with merge shown. Right-most panel shows micrographs with anti-OGA antibody for all three conditions. Scale bar, 10 μm. Inset scale bar, 5 μm. (B) Plot of the RL2 signal normalized to NUP96-GFP fluorescence in U2OS-CRISPR-NUP96-mEGFP cells treated as in (A). n>73 cells for each condition from three biological replicates. (C) As in B but with the anti-OGT antibodies to label fluorescence, n>33 cells from two biological replicates. (D) As in (C) but with anti-OGA antibodies to detect distribution of OGA, n>60 cells from two biological replicates. For (B-D) Kruskal-Wallis test with Dunn’s method for multiple comparisons performed. For (B): **, p=0.0012; ****, p<0.0001. For (C): ns, p=0.4161, ****, p<0.0001. For (D): ns, p=0.0910; *, p=0.0476. (E) Relative nuclear volume of iPSCs treated with the indicated drugs under normo-osmotic or hyper-osmotic (+ 400 mM sorbitol) conditions compared to the mean of the normo-osmotic DMSO condition (dashed line). Data represented as box and whiskers plot, box boundaries represent interquartile range, whiskers extend from minimum to maximum data point. Mixed-effects model with Sidak’s multiple comparison test performed. n=24-31 cells from one expansion experiment. ns, p=0.9395; *, p=0.0118. (F) Excess nuclear surface area of iPSCs treated with the indicated drugs under normo-osmotic or hyper-osmotic (+ 400 mM sorbitol) conditions. Data represented as box and whiskers plot, box boundaries represent interquartile range, whiskers extend from minimum to maximum data point. ns, p=0.6000; *, p=0.0202. n=24-31 cells from one expansion experiment. (G) Plot of the IR diameter assessed from pan-stained iPSCs subjected to the indicated drug treatment in hyper-osmotic conditions (+ 400 mM sorbitol) is plotted. Symbols denote individual cell means. Line and error bars are the overall mean ± SD. n=21,533 IR total in n=10 cells from one expansion experiment. Unpaired t-test. ns, p=0.0660

To provide insight into the mechanisms modulating nup GlcNAcylation upon hyper-osmotic treatment, we assessed the localization of the OGT and OGA enzymes. Under the hypo-osmotic condition we observed little change in the distribution of either enzyme, with both localizing throughout the cell with a modest concentration in the nucleus (Fig. 7A). Interestingly, however, in the hyper-osmotic condition we observed an obvious enrichment of OGT, but not OGA, along the nuclear periphery (Fig. 7A, C, D). Thus, under conditions in which the NPC is acutely constricted, the cell might respond by increasing nup GlcNAcylation, driven most likely by the recruitment of OGT to the NPC. Intuitively, this would counteract the constriction of NPC diameter by a decrease in the stringency of the diffusion barrier. To further test this model, we chose to mimic the increase in nup GlcNAcylation by pre-treating cells with TMG prior to the hyper-osmotic shock (Fig. S5A). In this case we observed that the loss of nuclear volume (Fig. 7E) and the excess of nuclear surface area (“wrinkling”)(Fig. 7F) arising from the hyper-osmotic shock are attenuated (Fig. 7E, F) as assessed by pan-ExM (Fig. S5B-G). Importantly, in this condition, the state of nup GlcNAcylation has no influence on the diameter of the IR as it remained constricted due to the loss of nuclear envelope tension (Fig. 7G). Together, these observations argue that cells rapidly adapt to NPC constriction by enhancing nup GlcNAcylation and decreasing the stringency of the diffusion barrier, supporting homeostasis.

## Discussion

The segregation of nucleoplasm and cytoplasm is determined by the integrity of the nuclear envelope membranes and by the selective permeability of NPCs. Here, we introduce the concept that the NPC diffusion barrier can be tuned by controlling the GlcNAcylation of FG-nups in response to both long-lived mechanical signals such as the stiffness of the extracellular environment and acute changes to nuclear envelope tension, such as those induced by external osmotic challenges. These new insights impinge on three important biological mechanisms: how cells tailor the function of NPCs to cell-type specific function, how cells execute mechanotransduction cascades, and how cells maintain the robustness of nucleocytoplasmic compartmentalization in the face of stress-induced changes to nuclear envelope tension.

### Mechanoresponsive nup GlcNAcylation ties diffusion barrier properties to the cell’s environment

Recent work provided evidence that either applying force on the nuclear envelope or the culturing of cells on “stiff” substrates weakens the NPC diffusion barrier, facilitating the translocation of macromolecules in the 27-41 kDa molecular weight range (Andreu *et al*., 2022). This “mechanosensitive nuclear transport” mechanism has been suggested to manifest through deformation of NPCs (Elosegui-Artola *et al*., 2017), perhaps NPC dilation (Matsuda and Mofrad, 2025). However, on first principles it is unclear if and how NPC dilation alters the sieving properties of the FG-nups. A key prediction emerging from our work is that cells in mechanically soft or stiff environments will also have correspondingly low or high levels of nup GlcNAc, respectively. We therefore argue that the interplay of NPC dilation state and nup GlcNAcylation together modulate the functional properties of the diffusion barrier, particularly given the evidence in prior studies (and reinforced by observations here) that GlcNAcylation can modulate the organization and dynamics of the FG network (Junod *et al*., 2025; Labokha *et al*., 2013; Yu *et al*., 2025). Nup GlcNAcylation will therefore be an important ingredient that must be incorporated into the emerging conceptual framework of mechanosensitive nuclear transport, which further highlights the need to define the mechanisms by which mechanical signals regulate nup GlcNAcylation.

Several additional challenges will need to be addressed to establish a molecular understanding of how mechanical signals are integrated with the GlcNAc machinery. First, as we find that NPC dilation and the GlcNAcylation state of nups are tightly correlated, disentangling how 1) nuclear envelope tension on NPC diameter and 2) direct impacts of coincident changes in nup GlcNAcylation on the NPC diffusion barrier together influence mechanosensitive nuclear transport will be challenging (see more below). Second, if and how LINC-complex dependent mechanotransduction, which has a modest impact on NPC diameter (Morgan *et al*., 2025), is integrated with the nup GlcNAcylation mechanism described here remains unclear. Of note SUN1 engages with NPCs, raising the possibility of direct force transmission to the NPC acting instead (or in addition to) nuclear envelope tension (Liu et al., 2007; Talamas and Hetzer, 2011).

Curiously, most of the stiffness-dependent changes in nup GlcNAcylation that we observe are independent of LINC complexes, suggesting that other pathways must translate mechanical signals to the GlcNAc machinery. Obvious candidates include actomyosin contractility, possibly regulated through MRTF (Infante et al., 2019) or Rho/ROCK (Hazawa et al., 2018; Nguyen et al., 2026), or mechanosensitive calcium release (Sarma and Yang, 2011). Mechanically-responsive kinases, including MAPK (Kosako et al., 2009), are also attractive candidates, particularly as kinases and OGT often compete for the same serine/threonine side chains (van der Laarse et al., 2018). Last, focal adhesion kinase (Lim et al., 2008), and indeed other focal adhesion components (Byron et al., 2022), are shuttling proteins that transit the NPC in a mechanosensitive manner (Cattaruzza et al., 2004; Nix and Beckerle, 1997). While the full nuclear function(s) of these factors remain to be defined, it is notable that TMG treatment, which increases NPC diameter, enhances the steady-state localization of zyxin in the nucleus, although this was previously ascribed to direct modification of the zyxin NES by GlcNAcylation (Zhao et al., 2022). With so many possibilities, there is likely to be a complex signaling network that is integrated in order to modulate the GlcNAc of the hundreds of potential nup GlcNAc sites (Ma *et al*., 2021), which may themselves be independently controlled by unique sets of inputs.

### Rapid changes in nup GlcNAcylation as a previously undiscovered homeostatic mechanism

While different mechanical states of the cell and targeted OGT/OGA engineering tools have revealed the steady-state coordination between NPC diameter and nup GlcNAcylation, we also provide remarkable evidence that cells execute a distinct response to a critical loss of nuclear envelope tension induced by hyper-osmotic shock. Here, OGT is rapidly recruited to NPCs to drive a condition that is not seen at steady-state: constricted NPCs with high nup GlcNAcylation. We posit that the recruitment of OGT serves two purposes: first, it promotes NPC dilation likely in an attempt to return the NPC to the preferred steady-state. Second, it makes the diffusion barrier more permissive, potentially facilitating the movement of osmolytes across NPCs to promote the re-equilibration of osmotic pressure across the nuclear envelope. This conclusion is supported by our observation that TMG treatment (high nup GlcNAcylation) promotes a less perturbed nuclear volume and nuclear envelope contour in response to hyper-osmotic shock despite little difference in NPC diameter. Further, this idea is consistent with at least one study that supports that the diameter of NPCs is wider in organisms like *Dictyostelium*, which may be an adaptation to facilitate water flow across NPCs to accommodate the rapidly changing osmotic environment relevant to this organism’s biology (Hoffmann *et al*., 2025). Importantly, although there is no GlcNAcylation in *Dictyostelium*, an analogous *O*-fucosylation modification likely occurs on nups (Tiwari et al., 2025), supporting the general conservation of this mechanism. It will be interesting to understand whether nup GlcNAcylation levels are higher in human cells within tissues that are exposed to fluctuating osmotic conditions such as the kidney. Indeed, genetic nup variants tied to steroid nephrotic syndrome already highlight that there may be important modifications made to NPCs in some kidney cells (Braun et al., 2018; Braun et al., 2016); if and how nup GlcNAcylation can explain these vulnerabilities awaits future work.

### The NPC diffusion barrier stringency in disease

Beyond fluctuating osmotics, nup GlcNAcylation is also likely to play cell-type specific roles that depend on the local extracellular matrix properties. We now understand that low nup GlcNAcylation is a property of cells in mechanically soft environments. Thus, neurons, which reside in soft tissue environments (∼1-2 kPa (Discher et al., 2005)), should have the strongest diffusion barriers; this prediction is borne out by the two neuron models that we have investigated here: dSH-SY5Y cells and iPSC-derived i^3^LMNs. Whether this is also observed *in vivo* is an important question and will be a priority for future work. That neurons have a stringent diffusion barrier is particularly interesting because of the ever-growing connections between the nuclear transport apparatus and neurodegenerative diseases (Chandra and Lusk, 2022; Coyne and Rothstein, 2022). We therefore wonder whether dysregulation of nup GlcNAcylation may be an early event in neurodegenerative disease pathomechanisms that could alter the distribution of specific macromolecules across the nuclear envelope. Key molecules that may be sensitive to a pathological weakening of the NPC diffusion barrier are proteins like the nuclear envelope ESCRT-III adapter CHMP7, whose passive nuclear entry is thought to be an early triggering event for ALS (Coyne *et al*., 2021); interestingly, the pathological entry of CHMP7 into the nucleus has been suggested to be modulated by the LINC complex component SUN1 (Baskerville et al., 2024), hinting at a potential sensitivity to mechanical inputs. Similarly, the leakage of TDP-43 out of the nucleus is thought to be a common, early event in ALS/FTD (Chen-Plotkin et al., 2010). Of particular note, both these factors have molecular weights from 40-50 kDa, at the edge of the regime that would be influenced by nup GlcNAcylation. Indeed, these new insights might also provide a rationale for the established links between the GlcNAcylation machinery and neuron health (Du et al., 2024; Wang et al., 2016; Zhu et al., 2014), although this possible connection will require further investigation.

### Integrating nuclear envelope tension, nup GlcNAcylation, and NPC dilation

That neurons and other cell types may tailor their nup GlcNAcylation profiles to promote cell type-specific (or condition-specific) function is compelling, but it remains unclear how each cell type establishes its nup GlcNAcylation “set point”. Our data support that mechanical cues are likely important players in dynamically establishing this set point, as the same cells plated on either soft or stiff substrates have low or high levels of nup GlcNAc, respectively. As substrate stiffness (Carley *et al*., 2021; Ihalainen et al., 2015), cell stretching (Nava et al., 2020), and osmotic shock (Enyedi *et al*., 2016), have previously been correlated with nuclear lamina and nuclear envelope tension, there may be a direct connection between the mechanical state of the nucleus with the GlcNAc machinery, or, alternatively, indirect mechanisms acting through one of the mechanotransduction mechanisms discussed above. Indeed, the recruitment of OGT to NPCs in response to loss of nuclear envelope tension represents additional evidence for the former. The physical properties of the diffusion barrier, which are likely weakened by GlcNAcylation, would act in opposition to nuclear envelope tension; the interplay of these two factors would therefore determine the NPC dilation state. Indeed, we establish that although modulating nup GlcNAcylation is sufficient to alter NPC dilation, a combination of high nuclear envelope tension and high nup GlcNAcylation is required to reach the largest NPC diameters even in the same mechanical condition.

There are two inter-related mechanisms that likely contribute to GlcNAc’s ability to promote NPC dilation: in one, the addition of GlcNAc may extend the unstructured, FG-rich domains, which may generate a steric pressure that effectively increases the volume in the central transport channel (Junod *et al*., 2025; Tan et al., 2018). Nup GlcNAcylation may also perturb direct FG-nup-NPC scaffold interactions that support NPC integrity (Onischenko et al., 2017). Alternatively, or in addition, the GlcNAc may disrupt cohesive FG-FG interactions that would be predicted to constrict the central channel (Labokha *et al*., 2013). In support of physical consequences of nup GlcNAcylation, a recent study leveraging time-resolved fluorescence anisotropy and fluorescently-labeled Nup98 revealed that OSMI-4 treatment (low nup GlcNAcylation) led to decreased segmental motions of the FG-repeat region while TMG treatment (high nup GlcNAcylation) moderately enhanced mobility *in vivo*, while *in vitro* elevating Nup98 GlcNAcylation promoted the maintenance of liquid-like behavior (Yu *et al*., 2025).

In terms of the ultimate functional consequences on the NPC diffusion barrier, the strong connection between nup GlcNAcylation and NPC dilation makes dissecting the contributions of the physical state of the FG network and NPC diameter changes challenging to disentangle. While we have furthered the evidence that engineering the elevated GlcNAcylation of the NPC is sufficient to enhance the diffusion of a reporter, this could be a direct effect on the behavior of the FG network and/or the effect of the coincident increase in NPC diameter. If the latter plays a role, it could be that a “thinning” of the barrier thickness in dilated NPCs and/or the collective increase in the total surface area of the NPC channel cross-section is responsible. Further progress into the detailed molecular mechanisms will require additional *in vivo* experimental tools, *in vitro* assays including use of DNA origami scaffolds to model NPC ring dilation and constriction, and *in silico* modeling.

## Materials and Methods

### Cell culture

HeLa cells (CCL-2; ATCC) were cultured in DMEM (11965092; Gibco) supplemented with 10% heat-inactivated fetal bovine serum (hiFBS; A5256801; Gibco), penicillin–streptomycin mix (pen/strep; 15140122; Gibco), and sodium pyruvate (11360070; Gibco). HEK293T cells (CRL-3216; ATCC) were cultured in DMEM supplemented with 10% hiFBS and pen/strep. A549 cells (CCL-185; ATCC) and *Sun1*^-/-^/*Sun2*^-/-^ double knock out (SUN1/2 dKO) A549 cells (Morgan *et al*., 2025) were cultured in DMEM/F12 (11320032; Gibco) supplemented with 10% hiFBS and pen/strep. HCT 116 cells (CRISPR edited HCT116 cell line with targeted insertion of tetracycline inducible *Oryza sativa* Transport Inhibitor Response 1 gene (Os-TIR1) at the AAVS1 locus (Natsume et al., 2016), were cultured in McCoy’s 5A modified medium (16600082, Gibco) supplemented with 10% hiFBS and pen/strep. SH-SY5Y cells (CRL-2266; ATCC) were cultured in DMEM/F12 (11320032; Gibco) supplemented with 10% hiFBS, pen/strep. U2OS cells CRISPR-edited to express a NUP96-mEGFP fusion from its endogenous locus (U2OS-CRISPR-NUP96-mEGFP, 300174; Cytion) was cultured in McCoy’s 5A modified medium (16600082; Gibco) supplemented with 10% hiFBS, pen/strep, 2 mM GlutaMAX (35050061; Gibco), sodium pyruvate, and MEM Non-Essential Amino Acids (NEAA; 11140050; Gibco). Human induced pluripotent stem cell (iPSC) lines WTC-mEGFP-Nup153-cl88 (AICS-0069-088; Coriell Institute) and WTC-mTagRFPT-LMNB1-cl62 (AICS-0034-062; Coriell Institute) were seeded on Geltrex (A1413302; Thermo Fisher Scientific) coated plates and cultured in mTESR1 media (85850; StemCell Technologies). All cells were maintained at 37°C with 5% CO_2_. To dissociate cells for passaging, cells were treated with 0.05% trypsin (25300054; Gibco), 0.5 mM EDTA (46-034-CI; Corning), or Accutase (SCR005; Sigma-Aldrich) in PBS.

### Cell line generation

WTC-mTagRFPT-LMNB1-cl62 iPSCs (AICS-0034; Allen Institute of Cell Science) were stably integrated with a doxycycline-inducible hNIL (NGN2-ISL1-LHX3) transgene cassette at the CLYBL safe harbor locus to allow differentiation into i^3^ lower motor neurons (i^3^LMNs) as described (Fernandopulle *et al*., 2018). Briefly, iPSCs were passaged as single cells with Accutase, seeded onto Matrigel-coated plates in complete mTeSR media supplemented with 10 μM ROCK inhibitor (SCM075; EMD Millipore) and transfected with Lipofectamine Stem (STEM00001; Invitrogen), Opti-MEM I (31985070; Gibco) and DNA plasmids (Table S1; CLYBL-(Ef1a-SBP-LNGFR-T2A-mApple)-(CAG-rtTA)-(TRE-hNIL), pZT-C13-L1, pZT-C13-R1, pCE-mp53DD). Enrichment for transfected cells was performed by magnetic streptavidin bead affinity (Dynabeads MyOne Streptavidin C1, 65001; Thermo Fisher Scientific). Following clonal isolation, successful line validation was confirmed via differentiation into i^3^LMNs.

### Neuronal differentiation

SH-SY5Y cells were differentiated into neuronal cultures using the retinoic-acid based method described in Shipley et al. (Shipley et al., 2016). Briefly, 200,000 cells were plated in a 6-well plate and cultured in “differentiation media #1” (DMEM/F12 with 2.5% hiFBS, pen/strep and 10 μM retinoic acid (R2625, Sigma-Aldrich)), which was replenished regularly. On day 7, cells were dissociated and replated. On day 8, the media was replaced with “differentiation media #2” (DMEM/F12 with 1% hiFBS, pen/strep and 10 μM retinoic acid). On day 10, cells were transferred onto ECM-coated dishes in “differentiation media #2” and then maintained in “differentiation media #3” (neurobasal media (21103-049; Gibco) with 1X B-27 (17504044; Gibco), 20 mM KCl, pen/strep, 2 mM GlutaMAX (35050061; Gibco), 50 ng/ml BDNF (450-02; Peprotech), 2 mM dibututyryl cyclin AMP (sc-201567; Santa Cruz Biotechnology) and 10 μM retinoic acid). Terminal differentiation was assessed by immunostaining for the presence of MAP2 and NeuN-positive cells and by the visualization of extensive and elongated neuritic projections. Typically, the cells were terminally differentiated at day 18 and were not used for experiments beyond day 30.

iPSCs with stably integrated doxycycline-inducible hNIL transgenes were differentiated into i^3^LMNs as described (Fernandopulle *et al*., 2018). Briefly, on day 1 cells were cultured in “induction media” (DMEM/F12 with HEPES, N2 supplement (17502048; Gibco), NEAA, GlutaMAX, 10 µM ROCK inhibitor, 2 µg/mL doxycycline (D3072; Sigma-Aldrich) and 0.2 µM Compound E (565790; Sigma-Aldrich)). On day 3, cells were passaged with Accutase and replated onto laminin and PLO coated plates in induction media with 10 μM ROCK inhibitor, 2 μg/ml doxycycline, 0.2 μM Compound E and 40 μM bromodeoxyuridine (BrdU, B9285; Sigma-Aldrich). On day 4, media was changed to “motor neuron culture medium” (Neurobasal medium with B-27 supplement, NEAA, N2 supplement, GlutaMAX with 1 μg/ml laminin). Typically, cells were terminally differentiated at day 7. After day 7, one-half culture medium changes were performed every 4 days and the cells were not used for experiments beyond day 15.

### Transfection and transduction

Plasmids were transfected into HeLa, A549 and HCT 116 cells at ∼70% confluency using lipofectamine LTX with PLUS reagent (15338100; Invitrogen) or into U2OS cells using lipofectamine 3000 (L3000150; Thermo Fisher Scientific) according to the manufacturer’s instructions. Plasmids were electroporated into neuronal dSH-SY5Y cells using a Lonza 4D-Nucleofector X Unit with (AAF-1003X; Lonza) in 20 μl 16-well strips and program CA-137. i^3^LMNs were transduced at 80% confluency with lentiviral particles at MOI of 0.5 and assayed 4-5 days post transduction. To perform siRNA knockdown of nup transcripts, HeLa cells were transfected with siRNAs against *NUP214* (L-011980-00-0005; Horizon Discovery), *NUP153* (L-005283-00-0005; Horizon Discovery), *NUP98* (L-013078-00-0005; Horizon Discovery), *NUP62* (4392420, Assay ID s24247; Ambion) or non-targeting siRNA control (D-001810-01-20; Horizon Discovery), at a final concentration of 60 nM using lipofectamine 3000 as per manufacturer’s protocol. Cell extracts were prepared 72 h post transfection and subjected to immunoblotting with respective antibodies.

### Lentivirus production

Using lipofectamine LTX with PLUS reagent according to the manufacturer’s instructions, lentiviral particles were generated by co-transfecting HEK293T cells at 60% confluency with lentiviral transfer vector (J81) containing the transgene (J81_EGFP, J81_EGFP1PrA or J81_EGFP2PrA) along with packaging and envelope plasmids psPAX2 and pMD2.G in a 1:1:1 molar ratio. After 16 h, media was replenished. Supernatants containing lentivirus were harvested 24 and 48 h later, pooled and clarified by centrifugation at 0.2 rcf for 5 min.

### Plasmids

All plasmids used in this study are listed in Table S1. The following plasmids were acquired from Addgene: pcDNA3.1(+)-myc-OGA(GS-544)C181 (Addgene plasmid # 193996; http://n2t.net/addgene:193996; RRID:Addgene_193996) and pcDNA3.1(+)-myc-OGA(GS-544)C181-D174N (Addgene plasmid # 193999; http://n2t.net/addgene:193999; RRID:Addgene_193999), both gifts from Christina Woo (Ge et al., 2023). CLYBL-(Ef1a-SBP-LNGFR-T2A-mApple)-(CAG-rtTA)-(TRE-hNIL)(gift from Michael Ward, Addgene plasmid # 105842; http://n2t.net/addgene:105842; RRID:Addgene_105842)(Fernandopulle *et al*., 2018); pZT-C13-L1 (Addgene plasmid # 62196; http://n2t.net/addgene:62196; RRID:Addgene_62196) and pZT-C13-R1 (Addgene plasmid # 62197; http://n2t.net/addgene:62197; RRID:Addgene_62197), gifts from Jizhong Zou (Cerbini et al., 2015); pCE-mp53D (gift from Shinya Yamanaka, Addgene plasmid # 41856; http://n2t.net/addgene:41856; RRID:Addgene_41856)(Okita et al., 2013); pMD2.G (Addgene plasmid # 12259; http://n2t.net/addgene:12259; RRID:Addgene_12259) and psPAX2 (Addgene plasmid # 12260; http://n2t.net/addgene:12260; RRID:Addgene_12260), gifts from Didier Trono. VB240319-1668ppt was generated using Vector Builder. pSC9_GBPOGA and pSC10_GBPOGA_D174N plasmids were generated by performing Gibson assemblies of PCR amplicons to replace the myc coding sequence with a Halo-GBP coding sequence in pcDNA3.1(+)-myc-OGA(GS-544)C181 or pcDNA3.1(+)-myc-OGA(GS-544)C181-D174N. To generate J81_EGFPxPrA plasmids, the EGFP-xPrA coding sequences were excised from IG062, IG024 and IG025 by Nhe1 and EcoR1 restriction digestion and ligated into J81 (gift from J. Rothstein, Johns Hopkins School of Medicine). To generate pSC11_mCh1PrA, the EGFP coding sequence was excised from IG062 by Age1 and BsrG1 restriction digestion and replaced with the mCherry gene. Plasmid sequences were verified by sequencing.

### GlcNAc-modifying drug treatments

To modulate whole cellular protein GlcNAcylation levels, cells were treated with either 10 μM OSMI-4 (HY-114361; MedChemExpress) to inhibit OGT or 3 μM TMG (13237; Cayman Chemical) to inhibit OGA for 24 h prior to performing experiments.

### Conditional modification of nup GlcNAc

To increase nup GlcNAc using a Halo-GBP-OGT fusion, U2OS-CRISPR-NUP96-mEGFP cells were transiently co-transfected with VB240319-1668ppt (containing the Halo-GBP-OGT coding sequence behind Tetracyline Response Element promoter) and VB010000-9369xhm (containing the Tet-controlled transactivator). After 24 h, Halo-GBP-OGT production was induced with doxycycline (D3072; Sigma-Aldrich) for 4 h. To decrease nup GlcNAc, U2OS-CRISPR-NUP96-mEGFP cells were transiently transfected with pSC9_GBPOGA (containing the Halo-GBP-OGA coding sequence behind CMV promoter). Twenty-four hours after transfection, OGA activity was induced with a 4 h treatment with 1 µM 4-hydroxy-tamoxifen (4-HT; S7827; Selleck chemicals), which triggers the removal of an intein engineered at cysteine 181 in the OGA gene (Ge *et al*., 2023). As a control, U2OS-CRISPR-NUP96-mEGFP cells were transiently transfected with pSC10_GBPOGA_D174N, which contains a Halo-GBP-OGA(D174N) encoding a catalytically inactive OGA.

### Osmotic shock

To expose cells to hypo-osmotic or hyper-osmotic conditions, the cell culture medium was replaced with either McCoy’s 5A modified medium (280-310 mOsm, for U2OS-CRISPR-NUP96-mEGFP cells) or mTeSR medium (330-350 mOsm, for the WTC-mEGFP-Nup153-cl88 iPSCs) diluted 1:1 with ddH_2_0 (resulting in 140-155 mOsm for McCoy’s 5A modified medium or 165-175 mOsm for mTeSR medium) or supplemented with respective medium containing 400 mM sorbitol (resulting in 680-710 mOsm for McCoy’s 5A modified medium or 730-750 mOsm for mTeSR medium) for time specified in figure legends, respectively.

### Immunoblotting

To generate protein extracts for immunoblotting, cells were scraped off cell culture dishes in RIPA buffer (50 mM Tris-HCl, 150 mM NaCl, 1% Triton X-100, 1% sodium deoxycholate, 0.1% SDS) supplemented with protease inhibitors (P8340; Sigma-Aldrich), phosphatase inhibitors (1862495; Thermo Fisher Scientific), and 40 μM TMG (13237; Cayman Chemical). Cell lysates were sonicated (microtip 4417; QSonica with 22-309783; Fisher Scientific) and clarified by centrifugation at 16,000 rcf for 10 min at 4°C. The protein concentration of the supernatants was determined using Bradford protein assays (5000006; Bio-Rad). 20 µg of total protein from cell extracts was mixed with an appropriate volume of 4X SDS-PAGE sample buffer (1610747; Bio-Rad), heat denatured and then resolved on 7.5% SDS-PAGE gels. The proteins were transferred to a 0.2 µm nitrocellulose membrane (1620112; Bio-Rad). Membranes were blocked in 5% non-fat milk (AB10109; AmericanBio) in tris-buffered saline-tween (TBST: 20 mM Tris-Cl, pH 7.5, 150 mM NaCl, 0.1% Tween 20) for 1 h at RT then incubated in primary antibodies. All antibodies were commercially sourced except for the antibody to NUP93, which we previously generated (Vishnoi et al., 2020). Information on all antibodies can be found in Table S2. Primary antibodies were diluted (Table S2) in 5% milk in TBST and incubated overnight at 4°C. After three washes in TBST, blots were incubated in secondary HRP-conjugated antibodies (Table S2) in TBST for 1 h. After extensive washing, ECL was visualized using SuperSignal West Femto Maximum Sensitivity ECL Substrate (34096; Thermo Fisher Scientific) on a VersaDoc imaging system (4000 MP; Bio-Rad) or ChemiDoc imaging system (12003153; Bio-Rad). Total protein loading was assessed using a Ponceau S Solution (P7170; Sigma-Aldrich).

### Immunofluorescence staining

Cells were fixed using 4% PFA (15710; Electron Microscopy Sciences) in PBS for 15 min, permeabilized with 0.2% Triton X-100 in PBS for 15 min, and blocked with PBS containing 5% BSA for 1 h at RT. Primary antibodies were diluted to concentrations indicated in Table S2 and used to label cells at 4°C overnight in a humidified chamber. After three washes in PBS containing 5% BSA, the cells were then incubated with secondary antibodies (Table S2), diluted in PBS containing 3% BSA for 1 h at RT. After three washes in PBS containing 3% BSA, cells were mounted onto microscope slides with Fluoromount-G mounting media (#17984-25; Electron Microscopy Sciences).

### Metabolic labeling and click-chemistry

U2OS-CRISPR-NUP96-mEGFP cells were metabolically labeled with 1 mM GlcNAz (MA30911; Carbosynth) in McCoy’s 5A modified medium for 48 h. Cells were fixed with 4% PFA for 15 min at RT. Subsequently, GlcNAz was labeled with CF640R-alkyne (92091; Biotium) in a Copper-catalyzed Azide-Alkyne Cycloaddition (CuAAC) click chemistry reaction (Yoo and Mitchison, 2021). Briefly,100 mM CuSO4 and 50 mM BTTAA were pre-mixed at a 1:5 ratio. The reaction cocktail was then prepared with 340 µl 100 mM sodium phosphate buffer, pH 7.0, 0.5 µl of 10 mM alkyne-CF640R, 110 µl of premixed CuSO4:BTTAA solution, 50 µl of 1 M sodium ascorbate. The cells were incubated with the reaction cocktail for 30-45 min at RT in the dark. Samples were then washed for 10 min twice in PBS containing 3% BSA and twice in PBS. The samples were then mounted onto slides for imaging with Fluoromount-G mounting media (#17984-25; Electron Microscopy Sciences).

### Pan-ExM and immunostaining of pan-ExM samples

Pan-ExM was performed as previously described (M’Saad and Bewersdorf, 2020; Morgan *et al*., 2025). Briefly, cells were fixed in 4% PFA in PBS for 1 h at RT, washed with PBS and post-fixed in 0.7% PFA and 1% acrylamide (AAm; 01697; Sigma-Aldrich) in PBS for 6 h at 37°C. The samples were then washed with PBS and embedded in first gelling solution (19% sodium acrylate (SA; S03880; Pfaltz & Bauer), 10% AAm, 0.1% N,N′-(1,2-dihydroxyethylene) bisacrylamide (DHEBA; 294381; Sigma-Aldrich), 0.25% ammonium persulfate (APS; 1610700; Bio-Rad), and 0.25% tetramethylethylenediamine (TEMED; T7024; Sigma-Aldrich) in PBS) within gelation chambers for 1.5 h at 37°C in a humidified container. Samples were denatured in 200 mM SDS (75746; Sigma-Aldrich], 50 mM Tris (T6066; Sigma-Aldrich), 50 mM sodium chloride (NaCl; 3627-07; JT Baker), pH 6.8) for 15 min at 37°C, then 1 h at 73°C and washed with PBS.

Gels were then stained with primary antibodies for 24 h and secondary antibodies (Table S2) for 12 h at 37°C. Both incubation steps were followed by three 15 min washes with PBST.

Gels were subsequently bathed in ddH_2_O until fully expanded and then incubated in second gelling solution (10% AAm, 0.05% DHEBA, 0.05% APS, and 0.05% TEMED) twice for 15 min each at on a rocking platform on ice. After removal of residual solution, gels were immobilized, placed in a humidified degassing chamber, and perfused with nitrogen gas for 10 min. The chamber was then sealed and incubated for 1.5 h at 37°C. Gels were then incubated in anchoring solution (2.8% PFA, 10% AAm) for 3 h at 37°C and washed in PBS for 30 min.

Next, gels were incubated in third gelling solution (19% SA, 10% AAm, 0.1% N,N′-methylenebis(acrylamide) (14602; Sigma-Aldrich), 0.05% APS, and 0.05% TEMED)) twice for 15 min each on a rocking platform on ice. After removal of residual solution, gels were immobilized, placed in a humidified degassing chamber, and perfused with nitrogen gas for 10 min. The chamber was then sealed and incubated for 1.5 h at 37°C. Gels were then incubated in 200 mM NaOH (7708-10; Macron) for 1 h and washed in PBS. The gels were then subjected to pan-staining with NHS ester CF568 and SYTOX Green staining (Table S2) and placed inddH_2_0 until fully expanded. Gels were mounted in #1.5H glass coverslip bottom µ-Slide 2 well dishes (Ibidi, 80287) and sealed with a two-component silicone (13001000; Picodent).

### Microscopy

The following fluorescence microscopes were used: (a) A Crest X-Light V3 Spinning Disk confocal system built on a Nikon Ti2-E Motorized Inverted Microscope with Perfect Focus System and four color D-LEDI widefield fluorescence, with a scientific complementary metal oxide semiconductor (sCMOS) camera (Teledyne Kinetix, 10 MP, 6.5 μm pixel), PlanApo 100X objective (1.45 NA). Tokai Hit Thermobox and stage top incubation system was to control temperature, humidity, and CO_2_. Images were acquired using the following filter sets: A Semrock FF01-391/477/549/639/741 multiband excitation filter was used with a Semrock FF421/491/567/659/776-DI01 multiband dichroic for Hoechst, mCherry, JF640, CF640R, and Alexa Fluor 647 imaging. Emission filters were FF02-438/24 (Hoechst), FF01-595/31 (mCherry), and Chroma ET665lp (JF640, CF640R, and Alexa Fluor 647). GFP, Alexa Fluor 488, and Alexa Fluor 594 were imaged using a Semrock FF409/493/596-DI02 dichroic with Chroma ET525/50m (GFP/Alexa Fluor 488) or Semrock FF01-647/57 (Alexa Fluor 594) emission filters. NIS-Elements was used to control imaging parameters. (b) A spinning disc confocal UltraVIEW VoX system built on an Olympus XI71 inverted microscope equipped with a PlanApo 60X objective (1.45 NA) and a Hammamatsu C9100-50 EMCCD controlled by the Volocity software (Improvision). GFP and mCherry images were acquired using 488 nm and 568 nm laser lines with appropriate filters (527 ± 23 nm filter and 594 ± 43 nm respectively). (c) An inverted Nikon Eclipse Ti2 microscope (Nikon Instruments) equipped with an Andor Dragonfly 600 spinning disk module (Oxford) and the Okolab enclosure for environmental control. The Dragonfly was used in confocal mode (40 µm pinhole disk) with a water-immersion PlanApo 40X (1.15 NA) and 60X objectives (1.2 NA) and imaged with a sCMOS camera (Sona 4BV6X, Andor Technologies). Dragonfly custom 405/488/561/640nm Quad Excitation Dichroic with 594/43 25mm diameter emission filter was used. Fusion software (Andor, version 2.4.0.22) was used to control imaging parameters.

### Bleaching experiments

Bleaching experiments to measure passive diffusion were performed on the UltraVIEW VoX, except in the case of cells cultured on hydrogels where to accommodate for substrate thickness and other limitations of the UltraVIEW VoX, fluorescence was acquired on the Nikon Eclipse Ti2 with Andor Dragonfly. Cells expressing GFP or mCherry-based reporters were grown on 35 mm glass bottom dishes (D35-10-1.5-N; CellVis) and incubated with medium lacking phenol-red prior to imaging. On the UltraVIEW VoX, the cytoplasmic fluorescence was bleached with 488 nm or 561 nm lasers. In the case of cells cultured on hydrogels, the Andor MicroPoint 4 photostimulation module was controlled by Andor IQ software to bleach the nucleus of selected cells with a 561 nm laser. In all cases, images were acquired at 1 s intervals for 10 s prior to bleach and for 300 s post bleach.

### Generation of PDMS hydrogels

Soft (∼3 kPa) or stiff (∼1.5 MPa) Polydimethylsiloxane (PDMS) substrates were prepared as previously described (Nassereddine et al., 2026; Seghir and Arscott, 2015). Briefly, PDMS hydrogels were generated by mixing base and cross linker components. Soft hydrogels (∼3 kPa) were generated mixing base and cross linker CY 52-276A/B (Dow Corning) at 1:1 ratio and stiff hydrogels (∼1.5 MPa) were generated by mixing base and cross linker Sylgard 184 (Dow Corning) at 1:10 ratio. The mixtures were degassed, added to 35 mm glass bottom dishes (D35-10-1.5-N; Cellvis) and spin coated at 1000 rpm for 60 s (Headway Research, PWM32). Both substrates were then cured at 70°C overnight. Before use, they were plasma cleaned (Gatan Solarus Advanced Plasma System (Model 950)) for 1 min, UV sterilized for 45 min, and then collagen coated for 1 h at 37°C.

### Substrate stiffness experiments

For assessing the RL2 levels when plated on substrates of different stiffness, U2OS or A549 cells were plated on either collagen-coated tissue culture plastic or collagen-coated 2 kPa Matrigen plates (PS100-COL-2-PK; Matrigen) for 48 h. For passive diffusion assays or immunofluorescence staining of cells on different substrates, U2OS or A549 cells were plated on collagen-coated PDMS substrates that were either soft (∼3 kPa) or stiff (∼1.5 MPa) for 48 h.

### Pan-ExM imaging

Pan-ExM imaging was performed on Nikon Eclipse Ti2 microscope with Andor Dragonfly 600 spinning disk module. SYTOX Green, CF568 succinimidyl ester, and ATTO647N were imaged with 488-, 561-, and 647-nm excitation, respectively. Entire cell volumes were acquired by performing z-stack tile scans using a 0.25-µm step size.

### Quantitative image analysis

Line profiles of fluorescence intensity of RL2 staining or NUP96-GFP were generated using the PlotProfile function in ImageJ/Fiji (Schindelin et al., 2012), using a 5-pixel line.

To quantify the fluorescence of GlcNAc labels at the nuclear periphery, the nuclear rim was first identified using custom scripts available at https://github.com/LusKingLab. In Figure 6E, a binary mask of each nucleus was generated from the Hoechst channel by thresholding the nuclear signal. The mask was sequentially eroded and dilated, and the resulting masks were subtracted to generate a ring-shaped region of interest (ROI) corresponding to the nuclear rim. This nuclear rim mask was then applied to both the GlcNAz (fluorescently labeled) and mAb414 fluorescence channel. The mean fluorescence intensity of GlcNAz was then normalized to that of the mAb414 signal. In Figures 3C, 4C and 4F the NUP96-GFP signal was thresholded to identify the nuclear rim. Here, the mean fluorescence intensity of the RL2 labeling at the nuclear rim was then normalized to that of NUP96-GFP intensity. In Figure 7, manual segmentation of the nuclear rim was performed to calculate the mean fluorescence intensity of RL2 labeling or the anti-OGT or anti-OGA labeling at the nuclear rim, normalized to the corresponding mean fluorescence intensity of NUP96-GFP.

To calculate passive diffusion rates, nuclear fluorescence was measured and background was subtracted. All measurement were normalized to the timepoint 20 s after bleaching (t=0 in plots). The rate of diffusion was calculated as the slope of a straight-line plotted from t=0 to t=40 s. For each field captured in timelapse imaging, we ensured that photobleaching was negligible by simultaneously monitoring and measuring nuclear fluorescence of an unbleached cell.

### Pan-ExM image segmentation and analysis

All 3D reconstruction, volume rendering and analysis of pan-ExM images were performed using Imaris versions 10.2-11.0 (Andor). Nuclei and NPC rings were segmented as previously described by LABKIT machine learning pixel classification (Morgan *et al*., 2025). Antibody signal was segmented using a background subtraction algorithm and the same manually determined threshold for all images.

Excess surface area (*ESA*) of segmented nuclei was calculated by comparing actual nuclear surface area (*A*) of each object to a sphere of equivalent volume (*A*_sphere_). The surface area of the isovolumetric sphere was defined as:

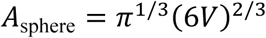

ESA was then computed as:

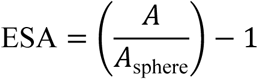

Expansion factors were determined by averaging peak-to-peak distances of line profiles drawn through centrioles and mitochondria in expanded samples using the Spots Intensity Profile Imaris XTension in MATLAB (The MathWorks). These values were divided by the previously determined dimensions of structures measured by EM to estimate the linear expansion factor.

### Data collection and statistical analysis

pan-ExM image segmentation data was compiled using Python (v3.12). All micrographs and immunoblots were analyzed in ImageJ/Fiji (Schindelin *et al*., 2012). All datasets from fluorescence imaging or immunoblots were collected and maintained in Microsoft Excel version 16.108. Statistical significance was assessed from data generated from independent experiments as indicated in legends and graphs were generated using GraphPad Prism version 11.0.1. Statistical tests are described in each figure legend. P<0.05 was deemed to be significant. Figures were assembled using Adobe illustrator version 30.3.

## Author contributions

S.C.: conceptualization, formal analysis, investigation, methodology, validation, visualization, writing – original draft, review & editing

K.M.: formal analysis, investigation, writing- review & editing

W.L.C.: formal analysis, software

C.P.L.: conceptualization, funding acquisition, project administration, resources, supervision, writing- original draft, review & editing

M.K.: conceptualization, funding acquisition, project administration, resources, supervision, writing- original draft, review & editing

## Acknowledgements

We thank Elisa C. Rodriguez for her invaluable support and help with cell culture, including differentiation and maintenance of motor neurons, Dr. Ivan Surovtsev for help with analysis pipelines, Krupa Hedge for assistance in maintenance of dSH-SY5Y cultures, Sandra D. Sandria and Valerie Horsley for assistance in preparation of hydrogels. We acknowledge the use of facilities and equipment at CINEMA (Cellular Imaging using New Microscopy Approaches) lab at Yale and especially thank Dr. Felix Rivera-Molina for his help with acquisition of live cell images for passive diffusion assay.

---

**Figure S1.**
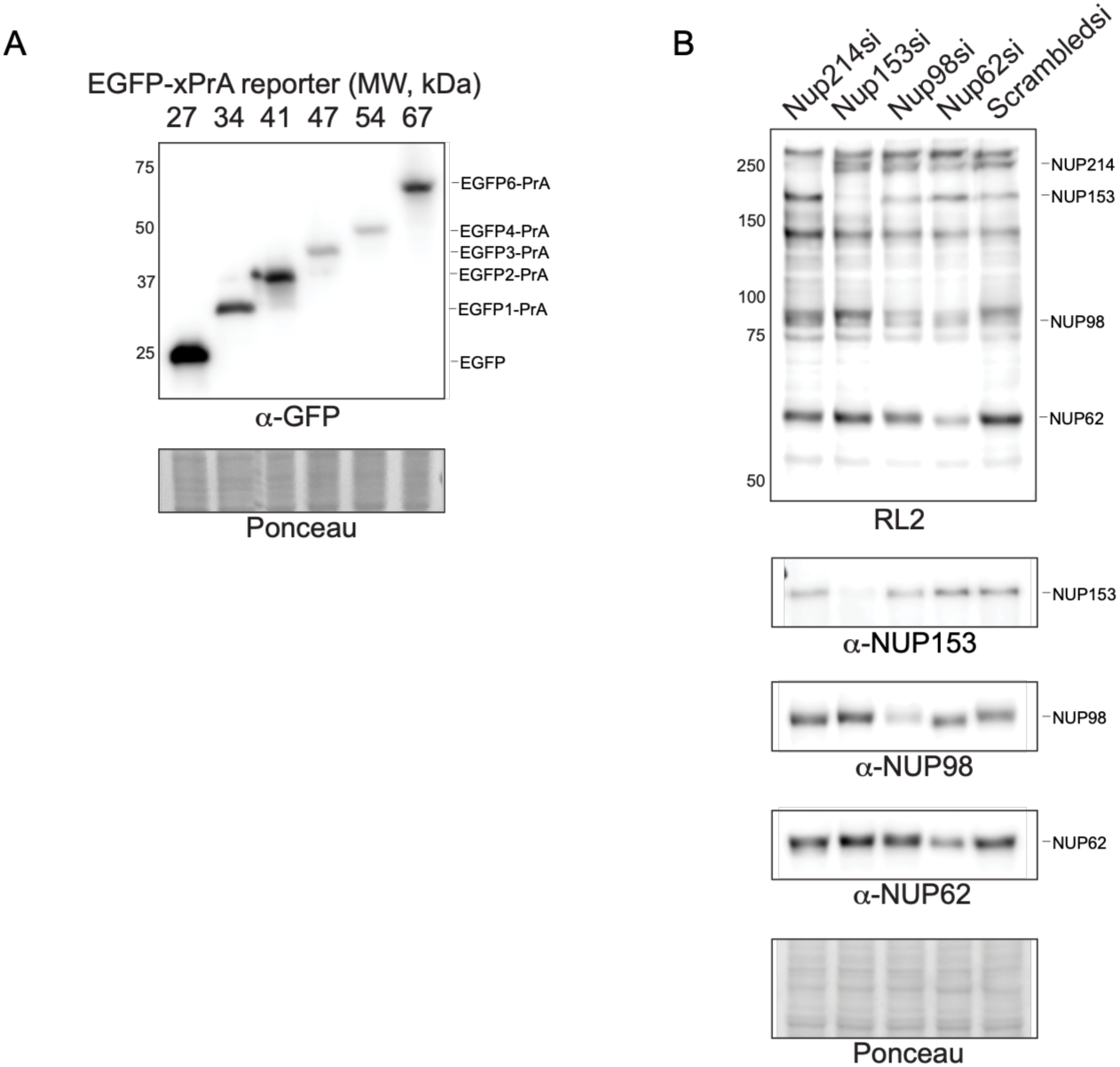
(A) Immunoblot (representative of two biological replicates) of protein lysates from HeLa cells expressing EGFP-xPrA reporters ranging from 27 kDa to 67 kDa, probed with GFP antibody. Numbers on the left indicate positions of molecular weight (MW) standards in kDa. To assess relative protein loading, a portion of the blot stained with Ponceau S is shown. (B) Confirmation of RL2-reactive nucleoporins by siRNA knockdown. Immunoblot (representative of 3 biological replicates) of protein lysates from HeLa cells, post 72 h treatment with siRNAs against *NUP214*, *NUP153*, *NUP98*, *NUP62* or a nontargeting control siRNA (Scrambled). GlcNAcylated proteins detected by RL2 antibody and nups with indicated antibodies. Numbers on the left indicate positions of molecular weight (MW) standards in kDa. To assess relative protein loading, a portion of the blot stained with Ponceau S is shown.

**Figure S2.**
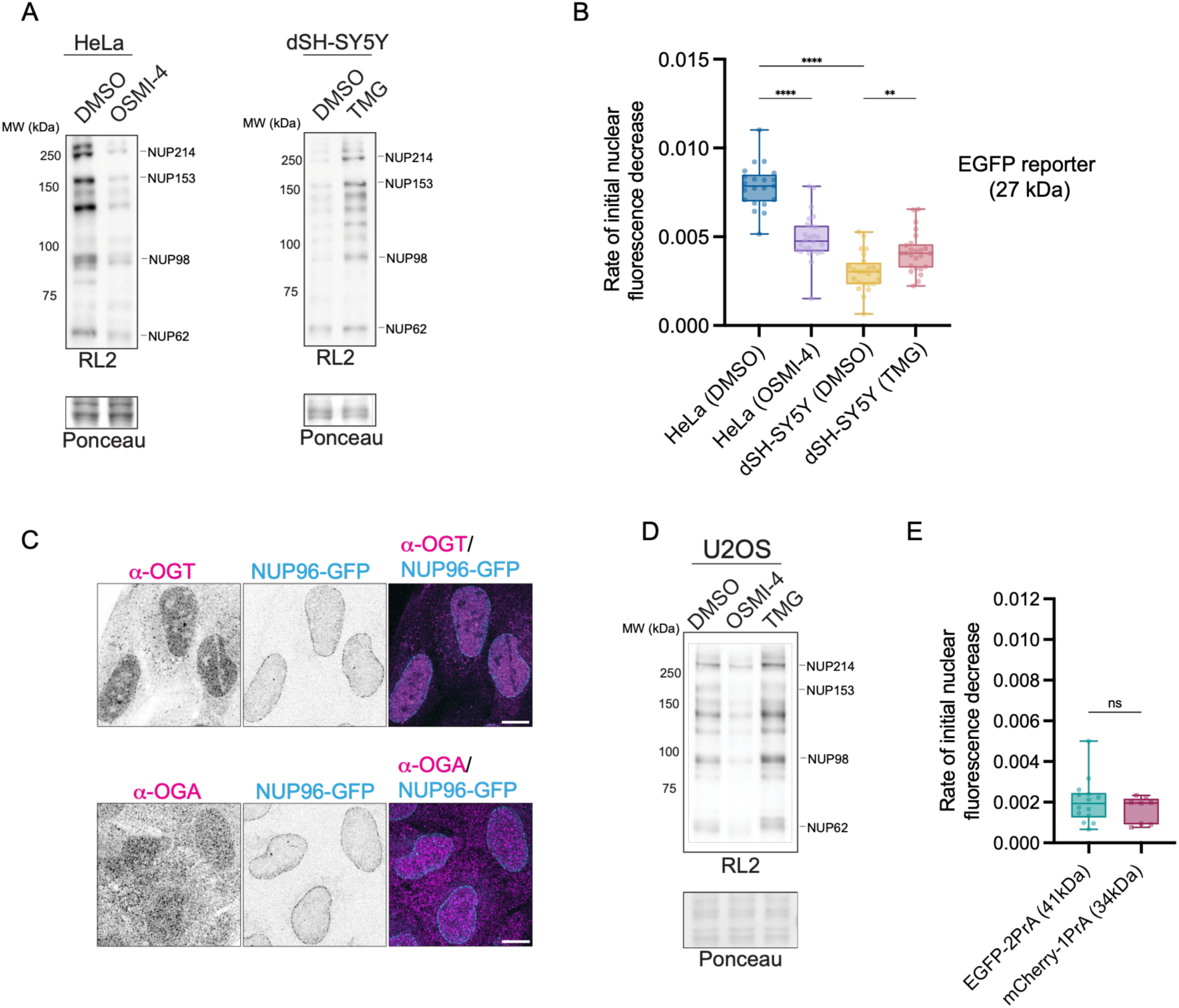
(A) Immunoblot (representative of two biological replicates) of protein extracts from HeLa cells treated with OSMI-4 or dSH-SY5Y cells treated with TMG (or DMSO as vehicle control) for 24 h and probed with the RL2 antibody. Numbers on the left indicate positions of molecular weight (MW) standards in kDa. Positions of nups as indicated. To assess relative protein loading, a portion of the blot stained with Ponceau S is shown. (B) Rate of nuclear fluorescence loss of 27 kDa EGFP reporter for cells treated as in (A). Plotted as in Fig. 1E. Data represented as box and whiskers plot, box boundaries represent interquartile range, whiskers extend from minimum to maximum data point, n>20 cells from two biological replicates, two-tailed Mann Whitney test performed. **, p=0.0054; ****, p<0.0001.(C) Confocal fluorescence micrographs of U2OS-CRISPR-NUP96-mEGFP cells stained with α-OGT or α-OGA. Images were acquired for NUP96-GFP, and anti-OGT or anti-OGA recognized by a secondary anti-rabbit antibody conjugated to Alexa fluor 647, scale bar 5 μm. (D) Immunoblot (representative of two biological replicates) of protein lysates from U2OS cells treated with OSMI-4 or TMG (as vehicle control) for 24 h as for panel (A). (E) Rate of nuclear fluorescence loss of indicated EGFP or mCherry-xPrA reporters in HeLa cells as in Figure 1E. Data represented as box and whiskers plot, box boundaries represent interquartile range, whiskers extend from minimum to maximum data point, n>8 cells from n=2 biological replicates, two-tailed Mann Whitney test performed. ns, p=0.3301.

**Figure S3.**
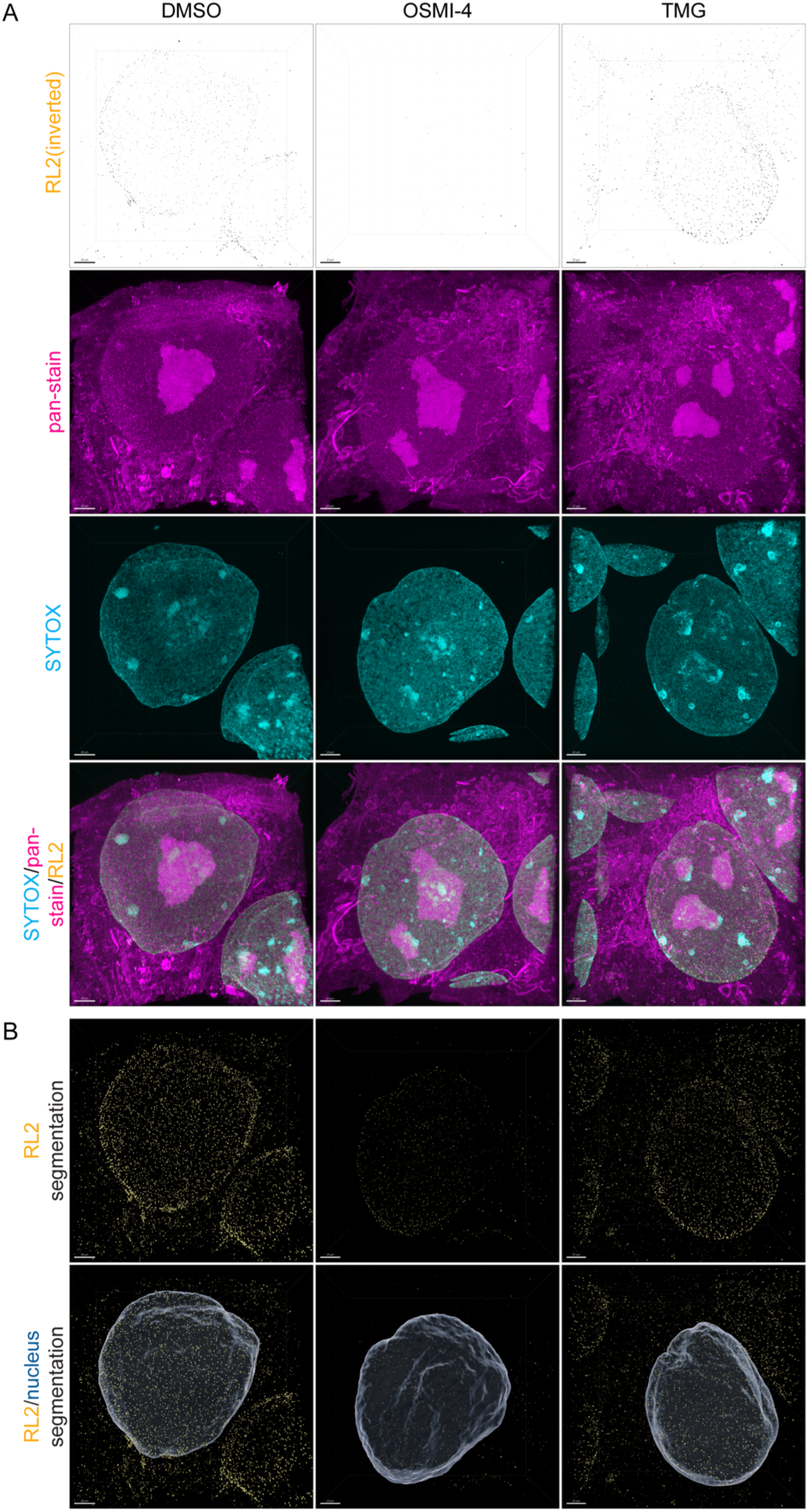
(A) A maximum intensity projection of a confocal fluorescence microscopy z-series of images comprising the entire volume of an iPSC prepared for pan-ExM and stained with RL2, a pan protein stain (NHS ester CF568) and SYTOX (for DNA). The cells were treated with DMSO, OSMI-4 or TMG for 24 h prior to fixation. The RL2, pan-stain, and SYTOX channels with merge are shown. Scale bar, 20 µm. (B) 3D renderings of RL2 and nuclei segmentation from cells in A. Scale bar 20 µm.

**Figure S4.**
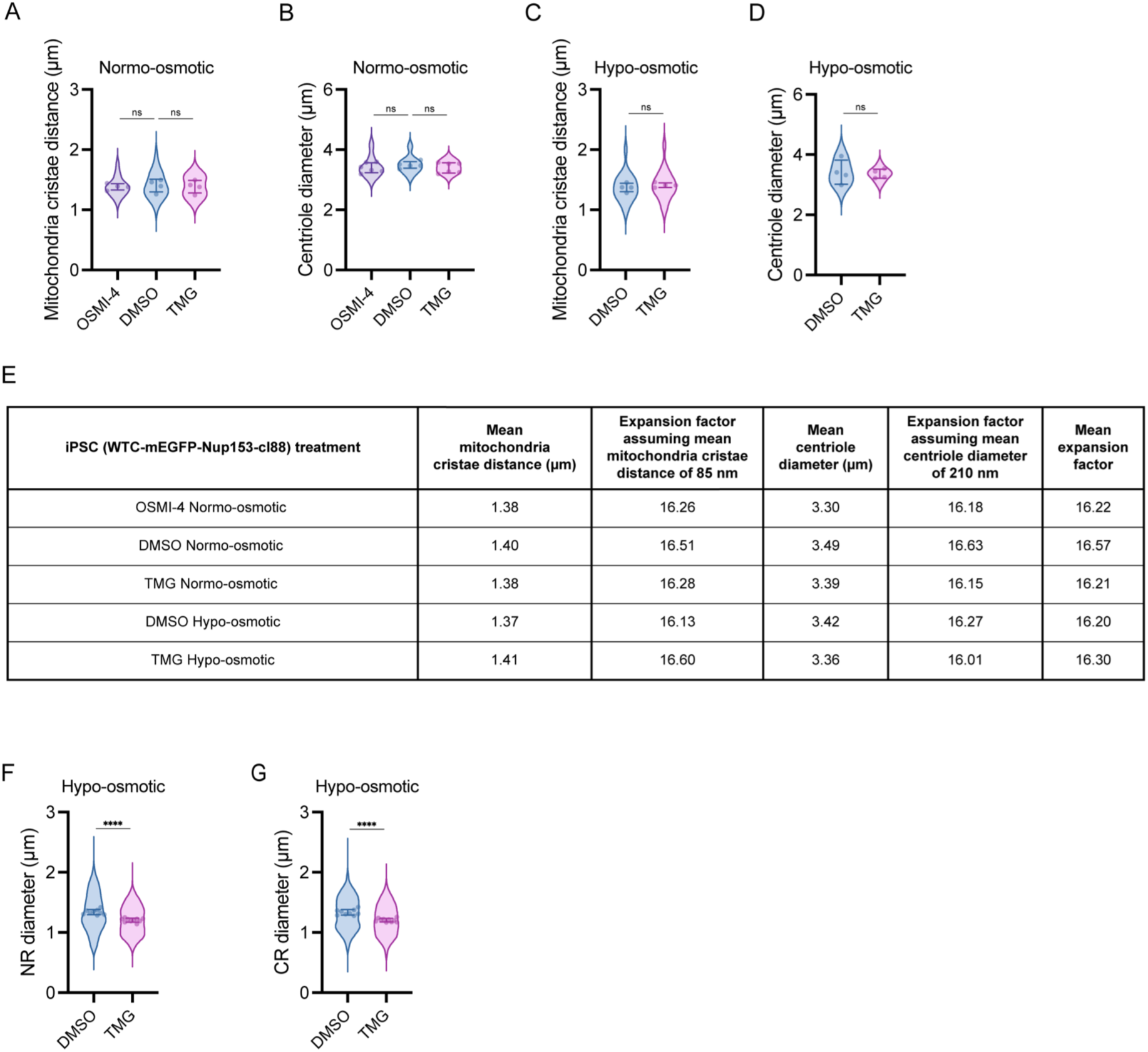
Quality control metrics for the robustness of pan-ExM across osmotic treatments. (A) Mitochondrial cristae distance in expanded iPSCs following the indicated drug treatments in normo-osmotic conditions. Symbols denote individual cell means. Line and error bars are the overall mean ± SD. n=5 mitochondria each in n=4 cells for each group from one expansion experiment. Ordinary one-way ANOVA with Dunnet’s multiple comparisons test performed. ns, p=0.9239; ns, p=0.9363. (B) Centriole diameter in expanded iPSCs following the indicated drug treatments in normo-osmotic conditions. n=10 centrioles each in n=5-6 cells for each group from one expansion experiment. ns, p=0.5005; ns, p=0.4720. (C) Mitochondrial cristae distance in expanded iPSCs following the indicated drug treatments in hypo-osmotic conditions (+ 400 mM sorbitol) plotted as in (A) n=5 mitochondria each in n=4 cells for each group from one expansion experiment. ns, p=0.5346; ns, p=0.5210. (D) Centriole diameter in expanded iPSCs following the indicated drug treatments in hypo-osmotic conditions (+ 400 mM sorbitol) plotted as in (B). n=8-10 centrioles each in n=4-5 cells for each group from one expansion experiment. ns, p=0.8554; ns, p=0.9393. (E) Summary table of post-expansion measurements of cellular structures and expansion factor calculations. (F-G) The diameter of the NPC nuclear ring (NR) (F) and cytoplasmic ring (CR) (G) following the indicated drug treatment in hypo-osmotic conditions (+ 400 mM sorbitol) is plotted. n=40,328 NR total in n=9-10 cells from one expansion experiment. **, p=0.0061; ****, p<0.0001; n=33,471 CR total in n=9-10 cells from one expansion experiment. **, p=0.0020; ****, p<0.0001.

**Figure S5.**
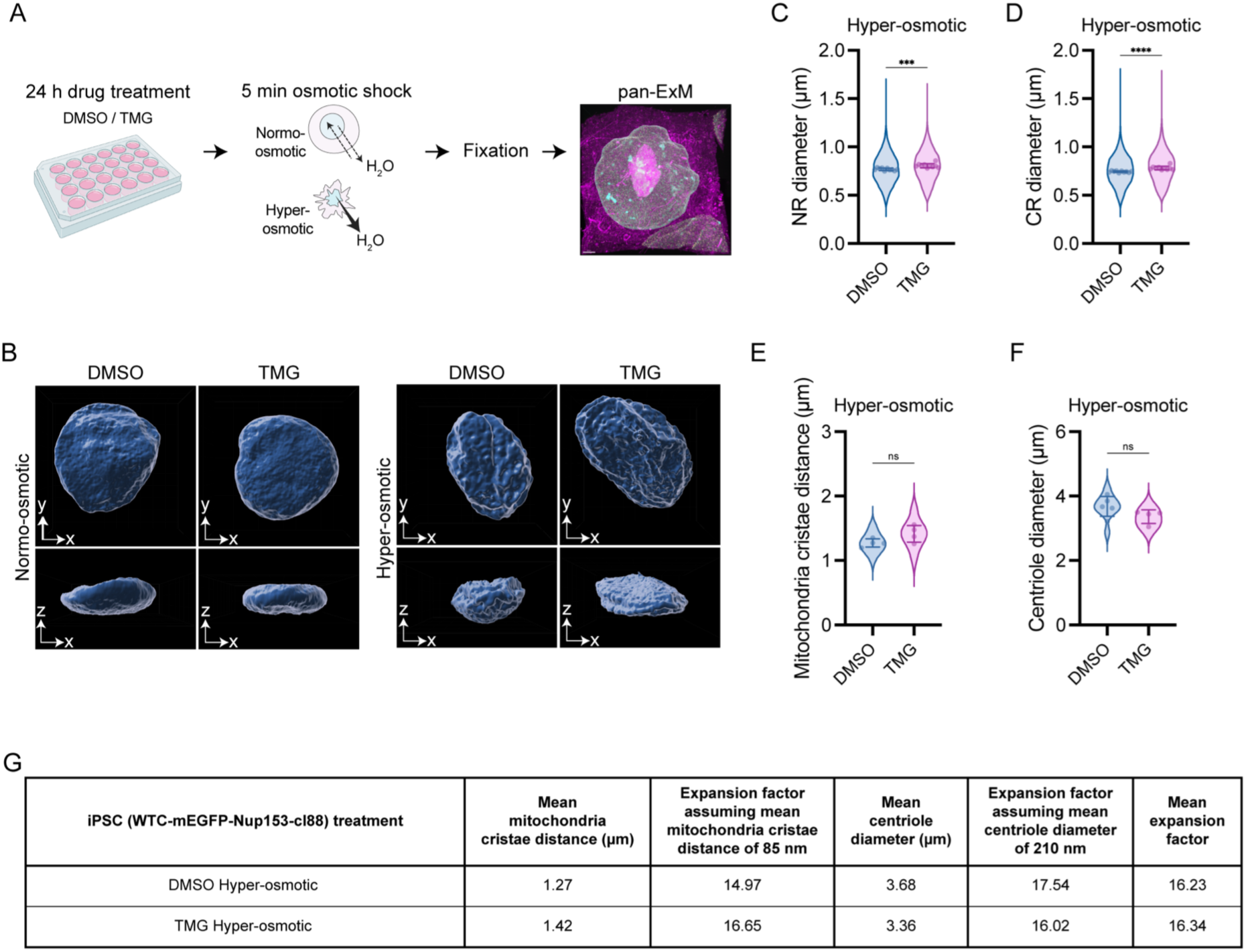
(A) Schematic of experimental workflow. iPSCs were treated with DMSO or TMG for 24 h, subjected to 5 min osmotic shock then immediately fixed for pan-ExM. (B) Representative 3D renderings of segmented nuclei from expanded iPSCs after the indicated drug treatment and in either normo- or hyper-osmotic (+ 400 mM sorbitol) conditions. (C) The diameter of the NPC NR following the indicated drug treatment in hyper-osmotic (+ 400 mM sorbitol) conditions is plotted. Symbols denote individual cell means. Line and error bars are the overall mean ± SD. n= 16,726 NR total in n=10 cells from one expansion experiment. Unpaired t-test performed. ***, p=0.0002. (D) As for (C) but for the NPC CR. n= 12,344 CR total in n=10 cells from one expansion experiment. ****, p<0.0001. (E) Mitochondrial cristae distance in expanded iPSCs following the indicated drug treatments in hyper-osmotic (+ 400 mM sorbitol) conditions. n=5 mitochondria each in n=4 cells from one expansion experiment. Unpaired t-test performed. ns, p=0.0945. (F) Centriole diameter in expanded iPSCs following the indicated drug treatment in hyper-osmotic (+ 400 mM sorbitol) conditions. n=9-10 centrioles each in n=4-5 cells from one expansion experiment. Unpaired t-test performed. ns, p=0.1239. (G) Summary of post-expansion measurements of cellular structures and expansion factor calculations for 5 min hyper-osmotic (+ 400 mM sorbitol) shock.

**Table S1:**
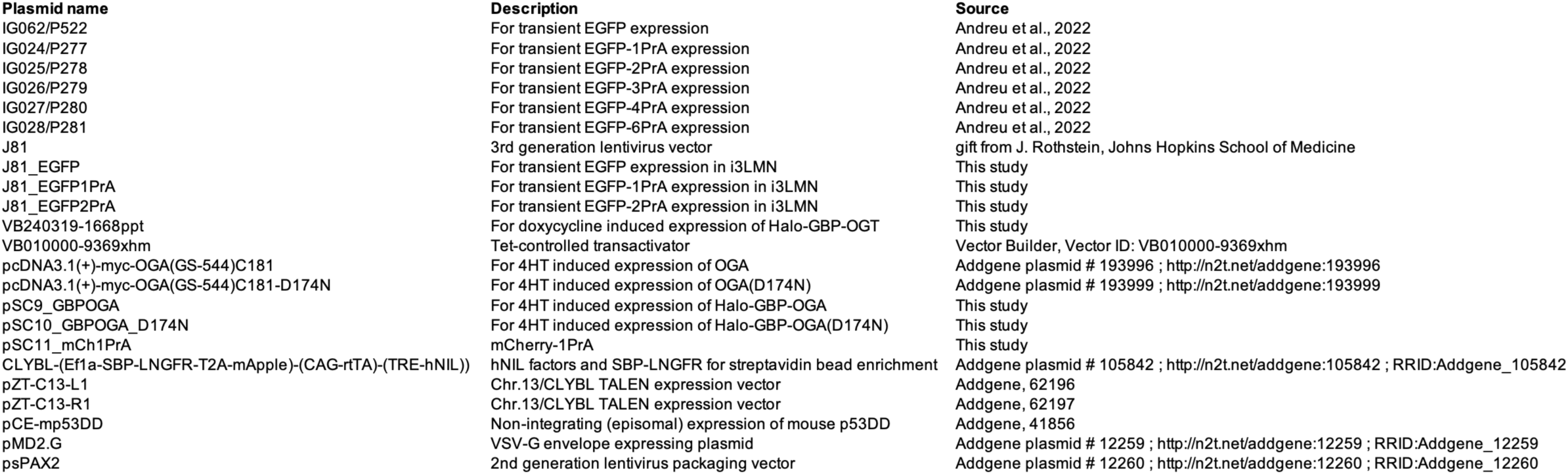
Plasmids.

**Table S2:** Reagents and Antibodies.

| **Antibody** | **Manufacturer/Source** | **Catalog number** | **Dilution used** |
| --- | --- | --- | --- |
| ATTO 647N (STED) Goat anti-Mouse | Active Motif | 15038 | 1:500 pan-ExM |
| Donkey anti-rabbit IgG, HRP-linked secondary antibody | Cytiva | NA934 | 1:10,000 WB |
| Goat anti rabbit IgG (H+L) secondary antibody, Alexa Fluor 647 | Invitrogen | A21245 | 1:1000 IF |
| Goat anti-Mouse IgG (H+L) HRP-linked secondary antibody | Invitrogen | 31430 | 1:10,000 WB |
| Goat anti-mouse IgG (H+L) secondary antibody, Alexa Fluor 488 | Invitrogen | A11029 | 1:1000 IF |
| Goat anti-mouse IgG1 secondary antibody, Alexa Fluor 594 | Invitrogen | A21125 | 1:1000 IF |
| Goat anti-rat IgG (H+L) HRP-linked secondary antibody | Invitrogen | 31470 | 1:10,000 WB |
| MGEA5/ OGA rabbit antibody | Abcam | ab124807 | 1:5000 WB, 1:100 IF |
| Nuclear Pore Complex proteins mAb414 antibody | Abcam | ab24609 | 1:250 IF |
| NUP62 mouse monoclonal antibody | BD Transduction Laboratories | BD610497 | 1:2000 WB |
| NUP93 rabbit polyclonal antibody | Vishnoi *et al.*, 2020 | N/A | 1:2000 WB |
| O-linked N-acetylglucosamine antibody (RL2) | Abcam | ab2739 | 1:2000 WB, 1:250 IF, 1:200 pan-ExM |
| OGT rabbit antibody | Abcam | ab177941 | 1:5000 WB, 1:100 IF |
| SUN2 rabbit monoclonal antibody | Abcam | ab124916 | 1:2000 WB |
| **Reagent** | **Manufacturer/Source** | **Catalog number** | **Dilution used** |
| CF568 Succinimidyl ester | Biotium | NC1542764 | 20 µg/mL pan-ExM |
| Hoechst 33342 | ThermoFisher Scientific | 62249 | 1:5000 IF |
| Janelia fluor 646 HaloTag ligand | Promega | GA1121 | 1:1000 IF |
| SYTOX Green | ThermoFisher Scientific | S7020 | 1:3000 pan-ExM |

